# Highly variable response of hard coral taxa to successive coral bleaching events (2019-2020) and rising ocean temperatures in Northeast Peninsular Malaysia

**DOI:** 10.1101/2021.11.16.468775

**Authors:** Sebastian Szereday, Affendi Yang Amri

## Abstract

Due to current greenhouse gas emissions, Malaysian coral reefs are predicted to experience severe annual coral bleaching events by 2043, threatening the survival of coral reefs within this century. However, there is no field data on how Malaysian coral reefs respond to successive events of coral bleaching. Despite the notion that many scleractinian taxa exhibit increased thermal tolerance over the last decade, it remains unresolved whether these changes are a result of ‘weeding out’ thermally susceptible species and actually ameliorate accelerating warming rates and increasing frequencies of heat disturbances. Moreover, complex interaction of environmental and biological factors that underlie differences in the bleaching response necessitate conducting studies at the within-reef scale (i.e., leeward shallow, windward shallow). Here, we studied two successive thermal stress events starting during the 2019 El Niño Southern Oscillation (ENSO), and determined bleaching trajectories of 29 hard coral taxa across fine spatio-temporal gradients in Northeast Peninsular Malaysia. Analysis of climate trajectories affirms accelerating warming rates (0.17°C per decade) and higher return-frequency of heat disturbance. Despite high annual maximum temperatures above the putative bleaching threshold (31.07°C and 31.74°C, respectively), accumulated thermal stress was low during both bleaching episodes (Degree Heating Weeks of 1.05°C-weeks and 0.18°C-weeks, respectively), suggesting widespread thermal sensitivity of hard coral taxa (55.21% and 26.63% bleaching incidence in 2019 and 2020, respectively). However, significant discrepancies between satellite and *in-situ* temperature data were found (0.63°C; SD±0.26). Bleaching severity was highly taxon-specific, varied across and within reef scales due to wind exposure and depth (e.g., less bleaching at shallow windward sites), and partially contrasted historical bleaching observations (e.g., *Acropora* and *Montipora* were less susceptible, *Cyphastrea, Echinopora, Goniastrea, Heliopora* and *Porites* were highly susceptible). While bleaching severity was higher in 2019, *Galaxea* and *Leptastrea* were bleaching more in 2020 despite lower heat stress, suggesting negative legacy effects of the 2019 bleaching event on these taxa. In conclusion, hard corals were subjected to more frequent heat stress during the last decade and remain highly vulnerable to marine heatwaves across all biophysical reef scales. Annual coral bleaching impacted all hard coral taxa and reduced thermal tolerance in at least two taxa.

## Introduction

The world’s oceans are the largest absorbers of energy derived from anthropogenic carbon-dioxide emissions (Pachauri et al., 2014), resulting in an increasing rapidity of ocean warming (Stocker et al., 2014). As the oceans warm, abnormally high sea surface temperatures (SST) induce global coral bleaching events with catastrophic consequences for the world’s coral reefs (Hughes et al., 2017). Coral bleaching refers to the whitening of coral tissue due to the disruption of its symbiotic relationship with its algae partner (Jaap, 1979). This symbiosis is essential for coral colony survival and is highly sensitive to thermal changes (Brown, 1997). Rapid ocean warming has thus been identified as the main threat to coral reefs worldwide (Hoegh-Guldberg, 2011), as the oceans have warmed 1°C over the past century (Hoegh-Guldberg, 1999), continuously exceeding the thermal maximum of scleractinian communities (Skirving et al., 2019; Van Hooidonk et al., 2020). Consequently, coral reefs are anticipated to decline globally by the end of the 21^st^ century and may not be capable of adapting to the projected increase in frequency and intensity of thermal events (Frieler et al., 2013), resulting in substantial shifts of foundation species within the coral reef assemblages, ultimately altering the baseline structure of these ecosystems (Loya et al., 2001; McClanahan et al., 2004; Hughes et al., 2018a). Such shifts are highlighted by recent studies that indicate higher resilience to heat disturbance of scleractinian communities that survived previous mass bleaching events (Van Woesik et al., 2012; Fox et al., 2019), and meta-analysis suggests large scale increases in thermal tolerance of hard corals populations (Sully et al., 2019). Nonetheless, thermo-tolerant populations presumably do not represent the primary assemblages of species that was found prior to frequent mass coral bleaching (Cannon et al., 2021).

As ocean warming rapidly increases (Cheng et al., 2021), annual severe bleaching (ASB) is predicted for 99% of the world’s coral reefs in this century (Van Hooidonk et al., 2013), underscoring a need to understand how consecutive episodes of coral bleaching will impact scleractinian community assemblage on fine spatial scales. In Peninsular Malaysia, ASB is predicted to occur by 2043 (Van Hooidonk et al., 2017), while regionally the first mass coral bleaching events were reported in 1998 and 2010, respectively (Kushairi, 1998; Tan and Heron, 2011). Annual coral bleaching episodes are still scarce, but may be adverse in impacts. The cumulative impact of annual coral bleaching may significantly alter scleractinian community assemblages and counteract thermal acclimation in hard corals (Grottoli et al., 2014). Acclimatization describes the genotype-independent response to physiological stress, observed on phenotypic-level and interpreting improvements in physiological stress response to changing and fluctuating environmental conditions (Edmunds and Gates, 2008). However, when adaptive responses are interpreted and contextualized as a response to a single environmental variable, the adaptive response is termed acclimation (Prosser 1991). Note, acclimation and acclimatization are not functional equivalents (Edmunds and Gates, 2008). Thus, rapidly rising ocean temperature represent a significant challenge to thermal acclimation of hard corals (Donner et al., 2005), and may undermine linear physiological adjustments, as heat stress exceed physiological thresholds more frequently, to a point where conceivable acclimation potentials to heat stress might be reduced and reversed (Schoepf et al., 2015). Conversely, the ecological footprint of large-scale heat disturbance events can also result in legacy effects that possibly precondition coral reefs at regional scales (Hughes et al., 2021), resulting in quick acclamatory response based on strengthening interactions of successive episodes (Hughes et al., 2019). As the time-intervals between such mass disturbance events are shrinking (Hughes et al., 2018b), it is no longer feasible to understand the cumulative impacts of coral bleaching by investigating isolated events, nor to assess ecosystem states without considering the legacy effects of previous disturbances (Johnstone et al., 2016). Reportedly, the shrinking time-intervals magnify the deleterious impacts by selectively pressuring more susceptible scleractinian taxa (Edmunds, 1994), resulting in winners and losers (van Woesik et al., 2011), and threaten ecological resilience and recovery (Hughes et al., 2021).

In 2010, changes in the hierarchy of bleaching susceptibility of scleractinian taxa were observed in southern Peninsular Malaysia, suggesting a capacity of certain taxa to adapt to rising SST based on previous thermal and bleaching exposure (Guest et al., 2012). Susceptibility to heat stress is highly variable among individual colonies and taxa (Marshall and Baird, 2000; Guest et al., 2016). On the level of coral reef community assemblages, heat stress susceptibility is underlined by the thermal history and variability experienced (Thompson and van Woesik, 2009; Thomas et al., 2018). Thus, adaptive responses that improve thermal tolerance are not uniformly confirmed in scleractinian communities (Hughes et al., 2017), due to differences in disturbance history and baseline temperature variability. Since the last studies reported on scleractinian bleaching susceptibility in southern Peninsular Malaysia in 2010, ocean warming has accelerated on a global scale (Skirving et al., 2019), culminating in the most prolonged and extensive pan-tropical coral bleaching event on record between 2014-2017 (Eakin et al., 2019). Thus, despite certain taxa showing adjustments in their thermal tolerance (*sensu* Guest et al., 2012), thermal stress and associated bleaching events are becoming more frequent, and it is ultimately unclear whether higher thermal stress frequencies and warming rates enhance or counteract acclamatory capacities (*sensu* thermal acclimation) of scleractinian taxa in Northeast Peninsular Malaysia and more broadly. Fine-scale assessments, spanning over repeated bleaching episodes, are important to understand scleractinian potential of thermal acclimation and to elucidate consequential shifts in scleractinian community assemblages. Furthermore, biological stress responses in hard corals are strongly correlated to environmental drivers (Fordyce et al., 2019), and are the result of a suite of complex biological and environmental interactions that underpin coral bleaching as a physiological stress response (Sugget and Smith 2019). Such abiotic-biophysical interactions include depth-dependent bleaching response (Bridge et al., 2014; Baird et al., 2018; Muñiz-Castillo and Arias-Gonzalez, 2021), and subsequent reef-wide effects of wind exposure (Page et al., 2019). Differences in environmental variability (i.e., light and wind) and subsequent high-frequency diel temperature variations of individual reef sites, influence heat tolerance on single reef scales substantially (Voolstra et al., 2020), and are possibly determining thermal tolerance beyond the levels that are manifested due to symbiont type and gene flow across reef sites (Oliver and Palumbi, 2011), ultimately highlighting the multifarious environmental influences on phenotypic stress response (*sensu* acclimatization) (Edmunds and Gates, 2008). Moreover, thermal tolerance acquired by surviving parent colonies in environmentally variable habitats are passed on to their offspring (Wong et al., 2021), and enable mechanisms of directional selection of heat tolerant corals through stress exposure, heritability, and reproduction. Hence, there is an urgent need to assess bleaching response of individual taxa in view of biophysical environmental variability within and across individual reef sites. Such understanding is crucial in order to uncover phenotypic variability across individual reefs (Palumbi et al., 2014; Safie et al., 2018). To date, few field studies have examined differences in bleaching response across fine biophysical scales and across successive events.

Due to rapid ocean warming, and under scenarios of continuous selection of bleaching tolerant species and subsequent reorganization of community assemblages, it is becoming important to derive thermal baselines from *in-situ* point measurements and in combination with field observations of bleaching response (McClanahan et al., 2007b; Heron et al., 2016a), to better comprehend the spatio-temporal variability of taxon-specific bleaching susceptibility at reef scale. Therefore, this study investigated the 1.) discrepancies between remotely sensed and *in-situ* temperature measurements to approximate reef-specific thermal bleaching thresholds and to analyse the accuracy of global web-based products (e.g., NOAA Coral Reef Watch). Secondly, 2.) the historical sea surface temperature trend and thermal anomalies above the upper threshold of scleractinian taxa were determined on island-wide scale, to highlight heat disturbance history and frequency, which conceivably propel or counteract adaptive legacy effects (Hughes et al., 2019; Hughes et al., 2021). Furthermore, we document the 3) taxon-specific bleaching susceptibility, recovery and mortality of scleractinian communities across the first annual sequence of coral bleaching in Northeast Peninsular Malaysia. During these successive thermal stress events in 2019 and 2020, 4) this study examined potentially disparate bleaching responses of scleractinian taxa. Lastly, to elucidate biophysical scales that potently induce acclamatory mechanisms that improve thermal tolerance in hard corals (Palumbi et al., 2014), we investigated 5) taxon-specific bleaching susceptibility across biophysical scales such as water depth and reef-specific wind exposure (i.e., leeward and windward), and partially determined the complex networking effect of wind, depth and diel temperature variations, which combined grossly span the physical gradient that underlines the ample environmental spectrum of hard coral resilience to heat stress.

## Materials and methods

### Study site

The present study investigated four coral reef sites around Pulau Lang Tengah (5°47’N, 102°53’E; Pulau=island) within the Terengganu Marine Park in Northeast Peninsular Malaysia (Figure 1). Two sites on the windward side (Batu Bulan (BB) and Tanjung Telunjuk (TT)) and two sites on the leeward side (House Reef (HR) and Secret Reef (SR)) were selected to explore whether biophysical factors such as depth and wind exposure of reef sites moderate fine-scale differences in the bleaching response of scleractinian taxa. Wind frequency direction was determined using web-based products^1^ and data references (Global Wind Atlas 3.0, 2020). Colony sample sizes across sites ranged between 323 and 663 colonies per site (mean 469.25 colonies/site). Noteworthy, leeward sites are chronically exposed to high sedimentation rates and sewage outflow from coastal tourism development and nearby resorts. In contrast, windward sites (particularly BB), are not exposed to immediate coastal development and are likely less impacted by sewage outflow from the nearest beach resorts.

**Figure 1.**
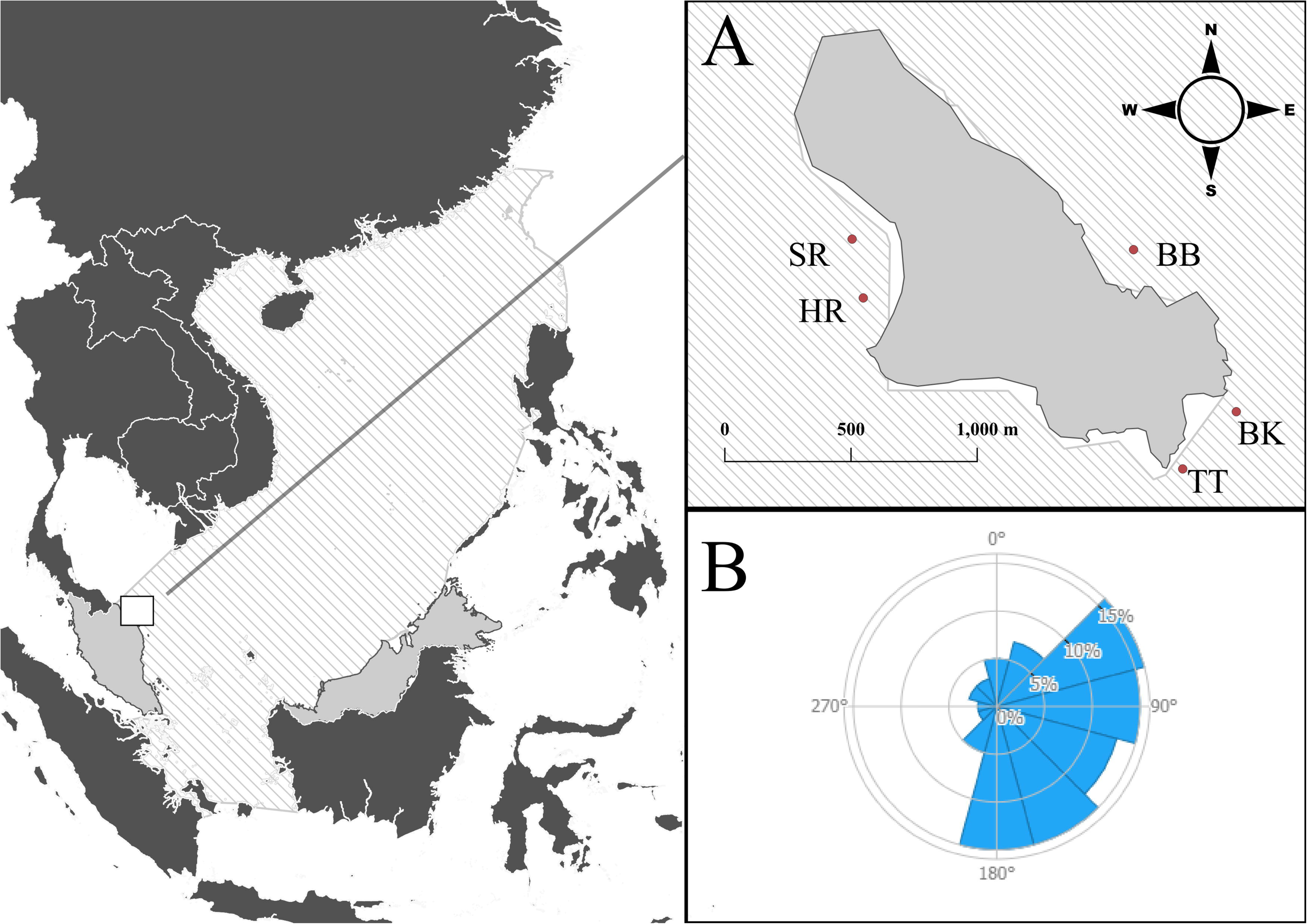
Location of Pulau Lang Tengah [(A), 5°47’N, 102°53’E] within the South China Sea (dashed area), showing survey sites [House Reef (HR), Secret Reef (SR), Tanjung Telunjuk (TJ) and Batu Bulan (BB)] and *in-situ* temperature recording stations [Batu Kucing (BK), House Reef, Tanjung Telunjuk]. The wind frequency rose (B) illustrates wind direction and frequency around Pulau Lang Tengah at island scale.

### Sea surface, *in-situ* and historical sea surface temperature analysis

Temperature records for Pulau Lang Tengah were sourced from remotely sensed sea surface temperature (SST) datasets through the National Oceanic and Atmospheric Administration’s (NOAA) next-generation operational daily global 5-km geostationary and polar-orbiting, blended night-only SST analysis, Coral Bleaching Monitoring Products (version 3.1, Coral Reef Watch, 2014), binned to the nearest 5 km^2^ satellite pixel (5°46’30.0”N, 102°52’30.0”E) and accessed through the ERDDAP website^2^. Note, that only nighttime SST are reported by satellites due to sunlight reflection on the sea surface and subsequent erroneous measurements. The accumulation of heat is expressed using the standard thermal stress metric “Degree Heating Weeks” (DHW) (Wellington et al., 2001). Additionally, we investigated the spatial extent of thermal stress in 2019 and 2020, by analyzing SST data for five additional locations across the east coast of Peninsular Malaysia, ranging from Pulau Sibu (∼420 km south, 2°13’14.7”N, 104°03’16.7”E), to Pulau Perhentian (∼35 km north, 5°56’02.5”N, 102°43’51.5”E).

Mean nightly sea surface temperature values between April 1^st^, 1985 and December 31^st^, 2020 were averaged for each year and linearly regressed in R Studio version 4.1.0 (R Core Team, 2021) to examine recurrent warming trends of our study location. In further steps, seasonality was eliminated by averaging SST for the six warmest months of the year (April– September). These months experience the least rainfall, cloud cover and storms, factors that significantly reduce bleaching severity (Skirving et al., 2014), and henceforth are referred to as the “bleaching season”, as coral bleaching exclusively occurs between April and September. Natural fluctuations leading to variability in SST was estimated by the standard deviation of the mean. Mean nightly sea surface temperatures were averaged for each decade, and past thermal disturbances were identified based on thermal anomalies that accumulated heat with values of DHW > 1°C-weeks (Liu et al., 2003; Kayanne, 2017). Coral bleaching occurs when SST exceed the climatologically warmest month of an area (MMM), by 1°C to create a coral bleaching hotspot (Glynn and D’Croz, 1990). Cumulatively, hotspots result in DHW (expressed in °C-weeks). DHW > 4°C-weeks induce ecologically significant bleaching, and DHW > 8°C-weeks results in ecologically significant mortality in scleractinian communities (Heron et al., 2016a; Skirving et al., 2019).

Five automated temperature monitors (HOBO 64K Pendant©, Onset Computer Corporation, USA) were deployed at three locations adjacent to transect sites at 8 meters’ depth. Data from TT and Batu Kucing (BK) reef was used to establish windward-site temperature regimes (Figure 1). *In-situ* measurements are not available for the first coral bleaching event in 2019, as monitor deployment was not conducted until the 11^th^ of September, 2019. Thus, the comparative analysis of *in-situ* monitors with remotely sensed data is based on data ranging from the 12^th^ of September 2019 – 30^th^ September of 2020 (19,635 measurements), using mean nightly temperature values (recorded each night between 00:00 – 06:00). A Pearson correlation analysis was used to examine data sets trends over the time period observed. Climatologically, the warmest month of the year in Pulau Lang Tengah is routinely in May, and the remotely sensed MMM is 29.94°C. However, the MMM based thresholds are putative and require repeated field observations of coral bleaching incidence and severity to confirm their accuracy. Since historical *in-situ* temperature data is not available, the MMM based on *in-situ* measurements is unknown. Therefore, for 2020, the mean monthly temperature difference between remotely sensed and *in-situ* measurements for May 2020 was calculated. The resulting difference was applied to the MMM of 29.94°C to establish an *in-situ* derived MMM of 30.51°C in order to calculate DHW values. This was repeated individually for each *in-situ* recording site to establish thresholds across biophysical reef scale, such as leeward and windward sites. To investigate site specific high-frequency variations in diel temperature, we calculated daily temperature ranges (DTR) between minimum and maximum temperatures, for each logging station for each day, and averaged DTR for the following time series: (DTR_Total_: September 2019 to September 2020), bleaching season (DTR_BS_: April to September 2020), peak summer season (DTR_SS_: May-July), monsoon season in boreal fall and winter (DTR_FW_: October-March), and for the 60 and 90 day period preceding the first bleaching observation in 2020 (June 20^th^, 2020), respectively (DTR_60_ and DTR_90_). Considering evident seasonal variability, we used a non-parametric Kruskal-Wallis test to determine seasonal dependence of DTR distributions (Salfie et al., 2018). To standardize DTR comparison across sites, only data from loggers logging at an interval of 90 minutes (HR, BK1, TT1) was used, excluding data obtained from two loggers recording at 15 minutes interval (BK2 and TT2). Lastly, the rapidity of heat stress onset and pre-stress exposure may function as a protective mechanism to mitigate subsequent bleaching (Ainsworth et al., 2019). To describe the accumulation of thermal stress prior to the first day of observed bleaching (6^th^ of June 2019, and 20^th^ of June 2020), we calculated the Degree Heating Days (DHDs) as the sum of all positive fluctuations of nightly SST above the MMM, following Maynard et al. (2008):

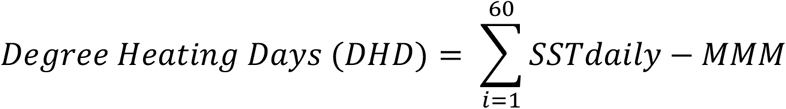

DHDs were calculated for the 60 and 90 day period prior to bleaching for each logging station and for remotely sensed data in in 2019 and 2020, respectively. The rate of heating (HR), which mathematically is the average rate at which DHDs accumulated throughout the 60 and 90 day period (Maynard et al., 2008), was calculated for all available temperature data:

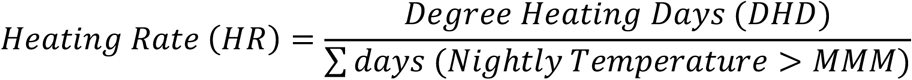

### Benthic community surveys

We hypothesize that rapid warming rates and more frequent thermal stress selectively influence scleractinian taxa, with conceivable impact on adaptive responses. Secondly, we investigated the response of scleractinian taxa to successive heat disturbances to better understand how such frequent heat stress will impact individual stress response in scleractinian taxa. As heat stress was less intensive in the second event, we particularly focus on taxa with equal or higher bleaching response in 2020 as compared to 2019. To differentiate between thermally tolerant and susceptible scleractinian taxa, a fine temporal analysis of the bleaching trajectory between June 2019 and June 2020 was conducted. Immediate bleaching response of scleractinian taxa was surveyed during thermal events in June 2019 and in June 2020, six and four weeks after SST exceeded the MMM by 1°C, respectively, whereby surveys were conducted three (in 2019) and seven (in 2020) days after the first signs of widespread bleaching were observed. Scleractinian community recovery from the 2019 event was surveyed in October 2019, 22 weeks post thermal stress onset. Surveys in April 2020, 50 weeks post thermal stress onset, were conducted to assess community recovery after the northeast monsoon season and before the next bleaching season in 2020. In this fashion, repeated and staggered assessment of the same scleractinian communities was carried out along permanently marked transect sites. A total of seven 10 × 1 m belt transects across the four sites (1,800 + coral colonies surveyed) were laid parallel to the shore on the shallower reef section between 5-7 m (mean depth 6 m), and on the deeper reef slope, between 10-14 m (mean depth 12 m), to account for differences in bleaching susceptibility at multiple depths. Each survey transect was marked with rebar stakes or PVC pipes at the start, middle (at 5m) and at the end point, and transect markers were number tagged. Additionally, transects were video recorded during each survey occasion for reference as to ensure the correct repositioning and redeployment of each transect tape during subsequent surveys. The numbers of healthy and bleached colonies encountered along the belt were recorded and identified to genus level following the Coral Finder 3.0 (Kelley, 2016). Only colonies whose colony centre was within the belt were recorded to avoid belt survey biases (Zvuloni et al., 2008). Bleaching severity categories were assigned for each coral colony: B1 (no bleaching), B2 (pale live and fluorescing colonies), B3 (≤33% of colony surface bleached), B4 (34% - 66%), B5 (67%-90%), B6 (>90% of colony surface bleached), where B5 and B6 are considered very severely bleached (Pratchett et al., 2013). Equally, dead corals were recorded to survey bleaching associated mortality and colony mortality was separated into these categories: M1 (≤33% of colony surface dead), M2 (34% - 66%), M3 (67%-90%), and M4 (>90%). Percentages of colony surface bleached or dead were visually estimated by the same observer across all survey occasions. Fluorescent bleaching was considered as very mild based on experimental findings (Bollati et al., 2020). The calcified octocoral *Heliopora coerulea* was included in the analysis, and the genus *Porites* was separated into *Porites* sp. 1 (massive and encrusting) and *Porites* sp. 2 (*Porites rus, Porites monticulosa* and similar species). A complete list of all anthozoan taxa observed bleached, including scleractinian taxa not recorded during belt surveys, is provided in Supplementary S1.

### Data and statistical analysis

Generalised Linear Models (GLM) (Nelder et al., 1972) with binomial distribution and a logit-link function were modelled to investigate coral bleaching susceptibilities (B1-B6) of selected scleractinian taxa (with at least ten observations, 11 taxa) in response to the effects of sampling time (June 2019, October 2019, April 2020, June 2020), wind exposure (leeward and windward), water depth (shallow or deep), taxon identity, and morphology. All explanatory variables were checked for multicollinearity prior to analysis. All GLM analyses were performed in R Studio version 4.1.0 (R Studio Team, 2021) using packages modEVA (Barbosa et al. 2014) and car (Fox and Weisberg, 2018). Following general linear model testing, we tested the interactive effects of depth and wind on site- and taxon-specific bleaching response using a Tukey’s post-hoc analysis to compare each interaction individually. Interaction were tested at fine biophysical scale (e.g., leeward shallow, leeward deep, windward shallow and windward deep) using emmeans-package v1.6.3 as a post-hoc test (Lenth, 2021). Finally, a Friedman test in R software was used to individually compare bleaching susceptibility (BR) across bleaching years for taxa that increased or maintained their BR in 2020 (four taxa), as well as for taxa who showed markedly higher resistance to thermal stress in 2020 as compared to 2019 (seven taxa).

To compare bleaching susceptibility of scleractinian taxa across annual events and environmental gradients (leeward, windward, shallow, deep), taxa with at least ten observations (n≥10) at each site (e.g., ten colonies at leeward and ten colonies at windward) were selected. The percentage of colonies per taxon in each group (B1-B6) was determined to calculate the standard Bleaching Response Index (BR) (McClanahan et al., 2007a), as to express bleaching susceptibility of each taxon at each site (leeward and windward), depth (shallow and deep), and sampling time (June and October 2019, June 2020):

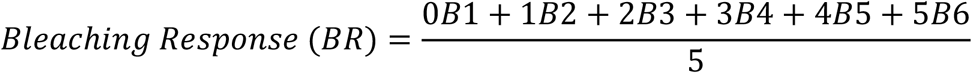

The BR Index is a normalized and weighted measure on a 0-100 scale. We focused our results on resistance to heat stress in order to express thermal tolerance of hard corals. Thus, we established the Bleaching Resistance Index as follows:

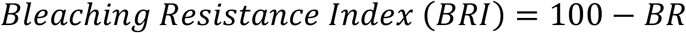

Furthermore, heat stress does not universally induce whole-colony mortality in all taxa, and partial mortality outweighs whole-colony mortality for numerous taxa (Baird and Marshall, 2002). Thus, heat stress induced mortality was separated from the Bleaching Index (BR) (as opposed to McClanahan et al., 2007a) and the Bleaching Induced Mortality Index (BIMI) was established instead, which accounts for partial- and whole-colony mortality as an immediate result of coral bleaching:

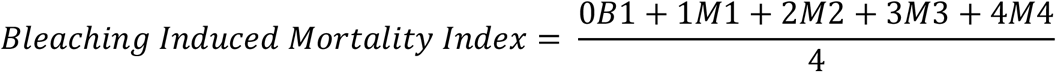

In order to account for notable differences in the relative density of taxa across environmental gradients (e.g., leeward and windward sites, shallow and deep transects), the site-specific Bleaching Susceptibility Index (McClanahan et al., 2007b) was applied, where *BR*_*i*_ is the taxon-specific bleaching response index following, *D*_*i*_ is the relative density of each taxon, and *N* the total amount of taxa:

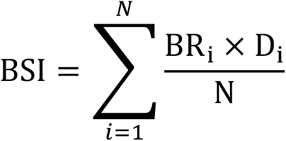

Summing each taxon’s BSI gave a site-specific bleaching index factoring in the relative density and total number of taxa.

## Results

### Sea surface, *in-situ* and historical sea surface temperature analysis

Linear regression analysis of annual mean SST (R^2^ =0.46; p<0.001) and mean SST of the bleaching season (R^2^=0.47; p<0.001) shows a warming trend since 1985 (Figure 2B and 2C), corresponding to an annual increase of 0.0166°C per year and 0.020°C per bleaching season. Starting in year 2001, minimum nightly SST started to steadily increase to above 27°C, and since 2015, the lower temperature range has not fallen below 27°C (Figure 2A). Mean SST increased rapidly since 2010, as SST during the bleaching seasons between 2010 and 2019 were 0.55±0.95°C (SD) higher as compared to mean bleaching season SST in 1985-1989, and 0.21±0.94°C higher than bleaching SST between 2000-2009 (Table 1). On average, 2020 and 2019 were the second (30.24°C±0.29°C) and fourth (30.12±0.59°C) warmest bleaching seasons since 1985, respectively. In addition to confirmed bleaching events in 1998, 2010, 2014, 2019, and 2020, six bleaching seasons between 1985-2019 were identified that reached accumulated heat stress values of DHW > 1°C-weeks. Two of these seasons occurred in 2013 and 2016, respectively, and reached similar or higher DHW values as in reported bleaching years in 1998, 2014, 2019 and 2020 (Table 1). Nine of the ten warmest years, and seven of the ten warmest bleaching seasons occurred between 2010 and 2020 (Supplementary S2).

**Table 1.** Mean decadal sea surface temperatures (°C) and temperature anomalies exceeding Degree Heating Weeks (DHW) 1°C-weeks are shown for Pulau Lang Tengah, Northeast Peninsular Malaysia. Years with observed and reported coral bleaching events are highlighted in grey and the source of *in-situ* coral bleaching observations is provided. Temperature measurements are sourced from National Oceanic and Atmospheric Association (NOAA), Coral Reef Watch (CRW) product, 2014, version 3.1.

| Decade | Mean annual SST °C (±SD) | Mean SST Bleaching Season °C (±SD) |
| --- | --- | --- |
| 1985-1989 | 28.86 (0.94) | 29.42 (0.95) |
| 1990-1999 | 28.88 (1.02) | 29.55 (1.02) |
| 2000-2009 | 29.05 (0.97) | 29.76 (0.94) |
| 2010-2019 | 29.30 (0.95) | 29.97 (0.95) |

| Year | Mean SST Bleaching Season<br>(°C) | DHW (°C-weeks) | Field observation |
| --- | --- | --- | --- |
| 1987 | 29.76 | 1.49 | - |
| 1990 | 29.35 | 1.14 | - |
| 1991 | 29.79 | 2.10 | - |
| **1998** | **30.13** | **0.34** | **Kushairi, 1998** |
| 2001 | 29.79 | 1.36 | - |
| **2010** | **30.38** | **5.16** | **Tan & Heron, 2011** |
| 2013 | 29.97 | 1.03 | - |
| **2014** | **30.10** | **1.60** | **Reef Check Malaysia, 2014** |
| 2016 | 30.09 | 1.01 | No bleaching (Szereday, personal observation) |
| **2019** | **30.12** | **1.05** | **This study** |
| **2020** | **30.24** | **0.00** | **This study** |

**Figure 2.**
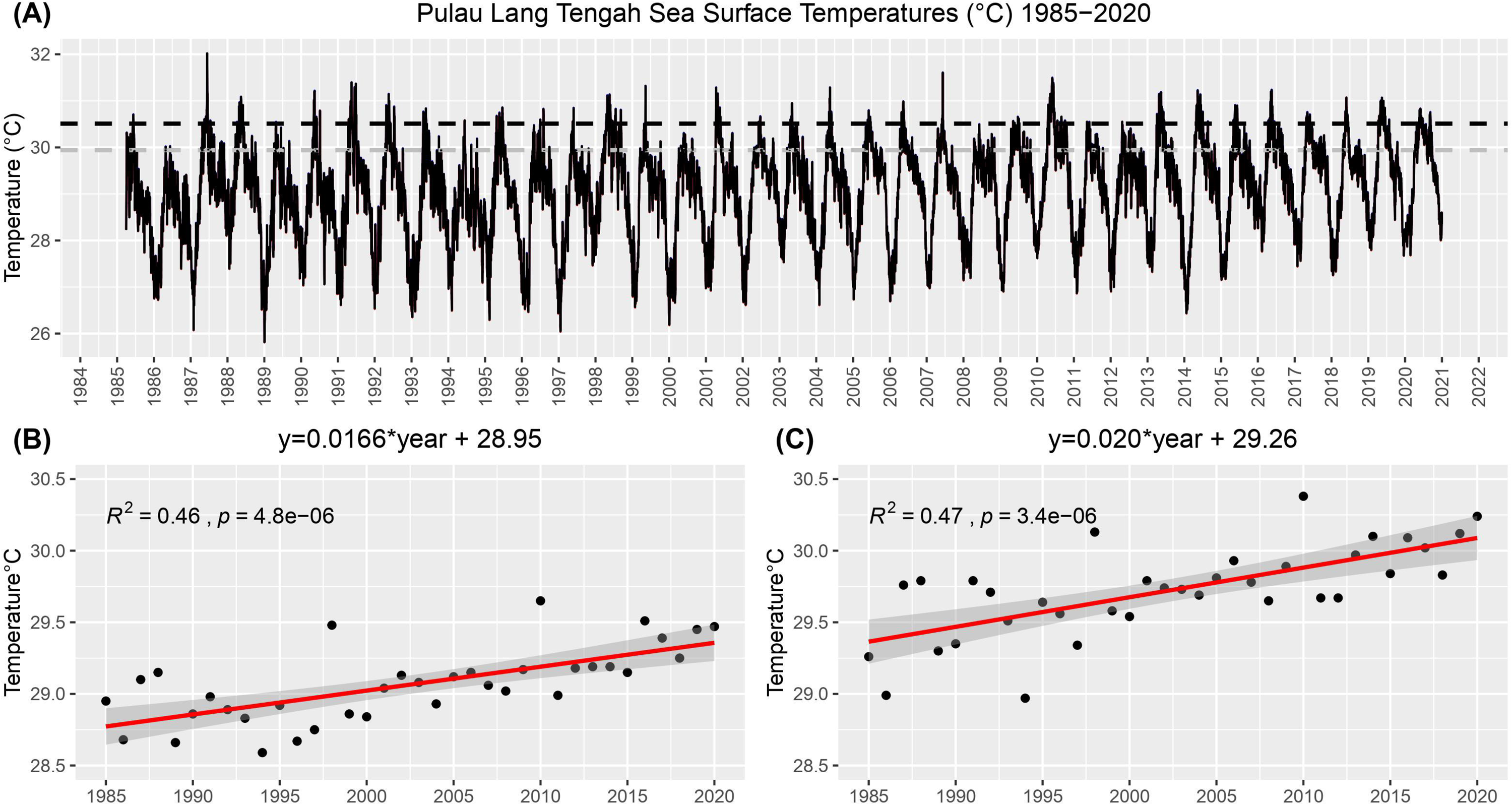
Nightly satellite measurements of sea surface temperatures (SST°C) around Pulau Lang Tengah between April 1985 and December 2020. The grey dotted line shows the maximum monthly mean (MMM=29.94°C) based on historical satellite measurements by the National Oceanic and Atmospheric Association (NOAA), Coral Reef Watch (CRW) product, version 3.1, and the black line presents the established *in-situ* MMM (30.51°C) after interpolating satellite data with *in-situ* measurements. Linear regressions of mean annual sea surface temperatures (B) and mean bleaching season sea surface temperatures (C) around Pulau Lang Tengah are shown. The regression line illustrates the warming trend since 1985, and the regression equation establishes average warming rates per year.

Heat stress in 2019 and 2020 was observed across the entire extent of the east coast of Peninsular Malaysia, with the highest values being recorded around Pulau Sibu (DHW = 4.61°C-weeks and 1.90°C–weeks, in 2019 and 2020 respectively) in Southeast Peninsular Malaysia (Supplementary S3). At our study location, *in-situ* measurements and remotely sensed measurements followed closely correlated patterns over the recording period from September 2019 to September 2020 (r=0.97, p<0.001), with a mean difference of 0.53±0.25°C. However, *in-situ* temperature measurements were 0.63±0.26°C warmer when analyzing temperature data for the bleaching season, whereas bleaching season means were less correlated (r=0.76, p<0.001) (Figure 3A). The rapidity of heat stress onset was markedly greater in 2019 for the 60 days period prior to bleaching (DHD_60_=41.33, HR_60_=0.69), compared to 2020 (DHD_60_=27.35, HR_60_=0.46), but not for the 90 days period (DHD_90_=24.20, HR_90_=0.27, and DHD_90_=20.29, HR_90_=0.23, in 2019 and 2020 respectively). Accumulated heat stress in 2019 measured remotely reached 1.05°C-weeks, and 0°C-weeks in 2020, not predicting levels of coral bleaching observed *in-situ* accurately in either year. Based on the MMM derived from *in-situ* measurements, DHW in 2020 reached 0.61°C-weeks in 2020 when pooling all *in-situ* measurements from all recording sites. At the time of surveying during the second bleaching event in 2020, heat stress accumulated at minor levels at 0.18°C-weeks (equivalent to two days of temperatures 1°C above the MMM). Nonetheless, observed bleaching levels did not correspond to the bleaching threshold as calculated by the *in-situ* derived MMM. Lastly, site specific analysis revealed fine-scale differences in temperature thresholds and heat stress (Figure 3, Supplementary S4), where average diel temperature variations (DTR) of each time series were significant more variable at leeward sites compared to windward sites (Supplementary S4), with the exception of the DTR_FW_ during the northeast monsoon season (Kruskal-Wallis test H=1.17, p=0.56).

**Figure 3.**
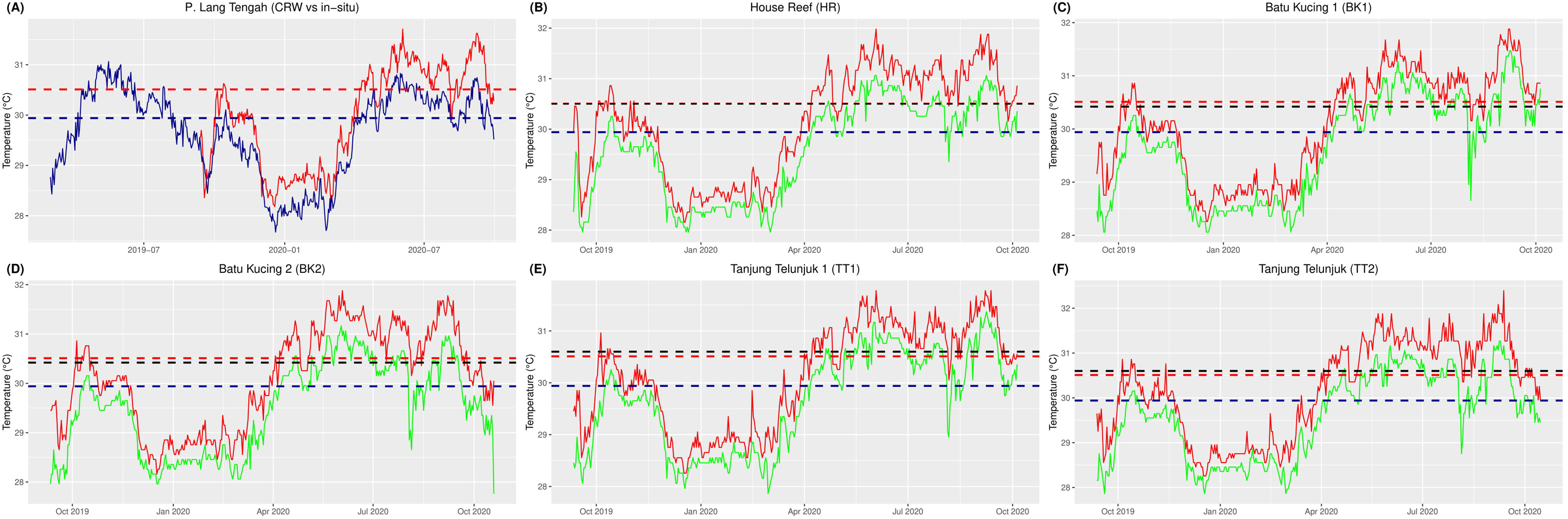
Nightly and diel temperature profiles at three sites around Pulau Lang Tengah, Malaysia. The blue dotted line presents the maximum monthly mean (MMM=29.94°C) based on the National Oceanic and Atmospheric Association (NOAA), Coral Reef Watch (CRW) product, version 3.1, and the red dotted line presents the *in-situ* MMM (30.51°C) after interpolating satellite data with *in-situ* measurements. The dotted black line shows the site-specific *in-situ* MMM after interpolation (MMM HR=30.50°C, MMM BK=30.42°C, MMM TT=30.60°C). **(A)** Comparison of satellite-derived (CRW) sea surface temperature (blue line) with average nightly *in-situ* point measurements (dark-red line) from three sites at 8 meters depth, recorded between September 2019 and September 2020, while the CRW time series starts in March, 2019. **(B-D)** Diel temperature profiles (°C) are shown as average daily (24h) maximum (red line) and average daily minimum (green line) temperatures for *in-situ* monitors logging every 30 minutes, and **(E-F)** for in-situ loggers recording every 15 minutes.

### Taxon-specific variability in annual bleaching response and recovery

Taxon-specific differences in bleaching response were chiefly pronounced. In June 2019, 55.21% of colonies surveyed (n=1,882) showed signs of bleaching, with 30.29% (n=570) of recorded colonies bleaching severely (Figure 4). Bleaching Resistance Index (BRI) in June 2019 was 64 out of 100, suggesting moderate tolerance to observed levels of heat stress. *Heliopora coerulea* showed the highest susceptibility in June 2019, followed by *Goniastrea, Cyphastrea, Pavona, Echinopora* and *Porites* sp. 1 (Table 2, Supplementary S5). In addition, *Porites* sp. 2 showed a severe bleaching response (BRI=66) but bleached less severely than *Porites* sp. 1. Of taxa historically considered most susceptible, *Acropora* (BRI=80) and *Montipora* (BRI=81) demonstrated less thermal susceptibility and higher tolerance, whereas *Pocillopora* bleached more strongly (BRI=60). Taxa with BRI > 90 were *Leptastrea, Symphyllia, Psammocora* and *Galaxea* (Table 2). Recovery rates were high in October 2019 as 91.68% of colonies surveyed (n=1,826) fully recovered and BI decreased significantly to 63.4 (BRI=97). The Bleaching Induced Mortality Index (BIMI) was universally low across all taxa (BIMI=1.71, n=1,826), whereas 1.10% of colonies (n=20) recorded full colony mortality, and 1.81% (n=33) experienced partial colony mortality. *Heliopora* showed the highest BIMI (11.36), recording 14.14% colonies with dead tissue, and 10.10% full colony mortality in October 2019. No bleaching was recorded in April 2020, and thus all taxa recovered fully from the 2019 bleaching event. In June 2020, moderate and frequent bleaching was recorded with a bleaching incidence of 26.63% of surveyed colonies (n=1,884). Bleaching intensity was significantly lower, as 6.26% of colonies bleached severely. General BI for all taxa was 231.6 (BRI=88), whereas *Heliopora coerulea* showed the highest bleaching response (BRI=59). Bleaching susceptibility moderately increased for two taxa in 2020 (*Galaxea* and *Leptastrea*) and *Symphyllia* showed similar bleaching response despite lower heat stress in 2020 (Table 2). In contrast, *Cyphastrea, Goniastrea* and *Echinopora* doubled their BRI in 2020, and *Favites, Fungia, Pavona* and *Porites* sp. 1 markedly increased their bleaching tolerance by at least 0.30. However, this change in bleaching susceptibility was only significant for *Fungia* (Friedman chi-squared = 4, df = 1, p-value = 0.0455) and *Porites* sp. 1 (Friedman chi-squared = 4, df = 1, p-value = 0.0455). Of all *Acropora* morphologies, only corymbose *Acropora* bleached in 2020. Morpho-taxon specific bleaching response is shown in the Supplementary S5.

**Table 2.** Bleaching Resistance Index (BRI) of the most abundant scleractinian taxa during two successive bleaching events in June 2019 and June 2020. The Bleaching Induced Mortality Index (BIMI) is shown for surveys conducted in October 2019.

| Genus | June 2019 |  | October 2019 |  |  | June 20 |  |
| --- | --- | --- | --- | --- | --- | --- | --- |
|  | n | BRI | n | BRI | BIMI | n | BRI |
| *Acropora* | 220 | 80 | 158 | 100 | 0.16 | 171 | 98 |
| *Cyphastrea* | 19 | 40 | 25 | 100 | 0.00 | 22 | 92 |
| *Echinopora* | 34 | 44 | 25 | 98 | 2.00 | 24 | 96 |
| *Favia* | 71 | 82 | 96 | 95 | 1.30 | 114 | 96 |
| *Favites* | 38 | 57 | 35 | 100 | 0.71 | 35 | 93 |
| *Fungia* | 115 | 61 | 120 | 89 | 0.83 | 137 | 91 |
| *Galaxea* | 96 | 98 | 110 | 100 | 0.00 | 101 | 95 |
| *Goniastrea* | 52 | 34 | 61 | 97 | 0.00 | 44 | 85 |
| *Heliopora* | 122 | 18 | 99 | 76 | 11.36 | 92 | 59 |
| *Leptastrea* | 69 | 93 | 72 | 99 | 0.00 | 80 | 92 |
| *Montipora* | 51 | 81 | 40 | 99 | 0.63 | 44 | 96 |
| *Pavona* | 144 | 41 | 125 | 95 | 3.40 | 130 | 74 |
| *Platygyra* | 68 | 73 | 95 | 94 | 1.58 | 103 | 81 |
| *Pocillopora* | 29 | 60 | 31 | 96 | 4.84 | 38 | 85 |
| *Porites* sp. 1 | 202 | 47 | 232 | 100 | 2.69 | 262 | 82 |
| *Porites* sp. 2 | 447 | 66 | 384 | 100 | 0.78 | 354 | 88 |
| *Psammocora* | 42 | 98 | 45 | 100 | 0.00 | 48 | 100 |
| *Symphyllia* | 26 | 94 | 31 | 100 | 3.23 | 34 | 96 |
| **Total** | **1,845** | **0.63** | **1,784** | **97** | **1.71** | **1,833** | **87** |

**Figure 4.**
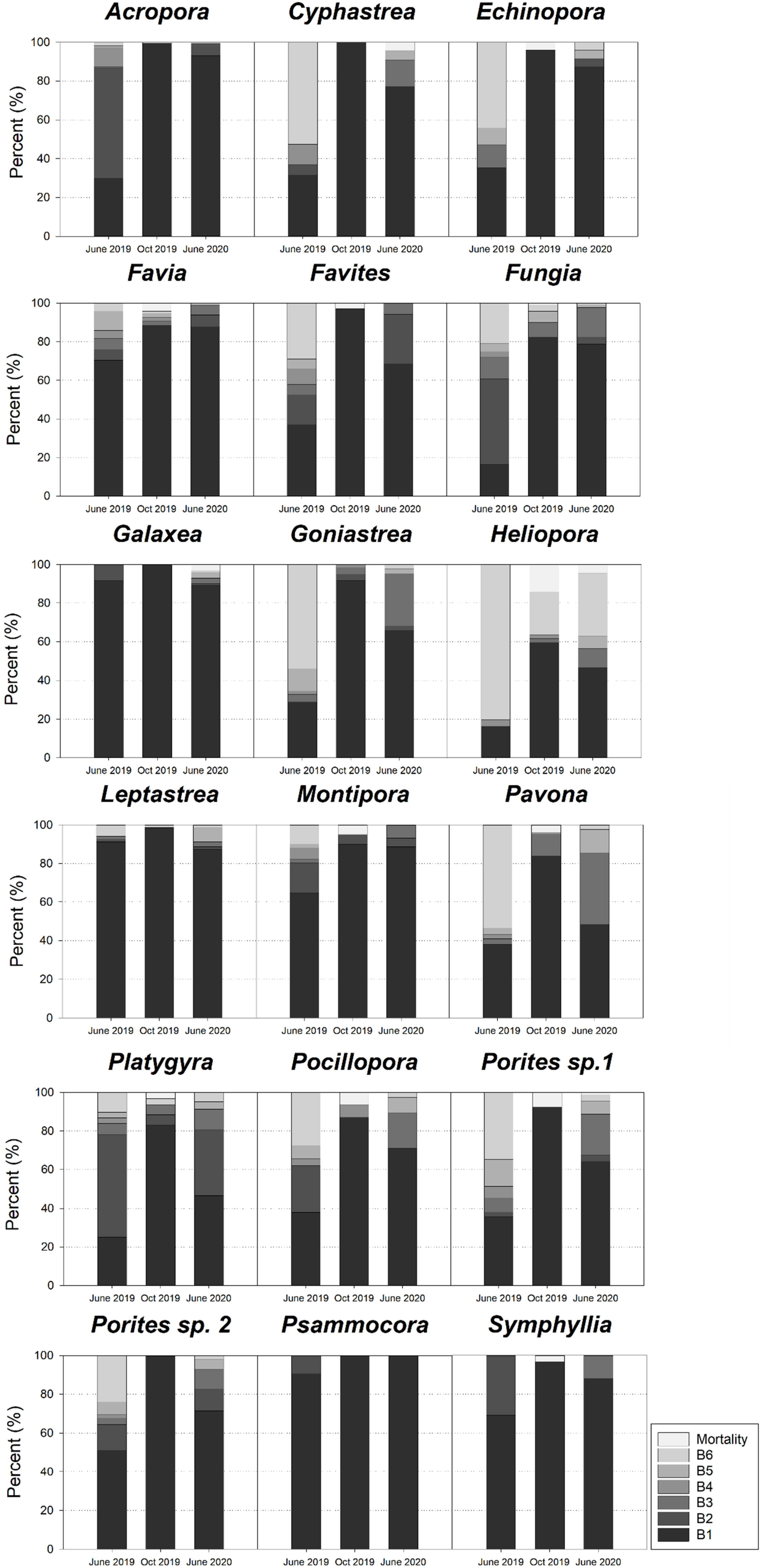
Percentage of healthy, fully and partially bleached colonies of scleractinian taxa with at least ten observations during survey occasion in June 2019, October 2019 and June 2020. In October 2019, percentage of bleaching induced mortality is shown, combining complete and partial-colony mortality.

### Bleaching response on biophysical reef scale

Bleaching incidence in 2019 was higher at leeward site (70.82%) compared to windward sites (44.85%). Generalized linear models tested seven taxa (n≥10 at both sites) for significant differences in bleaching response across leeward and windward sites. *Acropora, Echinopora, Fungia, Pavona* and *Porites* sp. 2 bleached significantly less at windward sites in 2019 (Table 3), whereas in 2020, *Fungia* and *Porites* sp. 1 bleached significantly less at windward sites (Table 3). Pooling data from both years and repeating the analysis revealed that *Acropora, Fungia* and *Pavona*, bleached significantly less at windward sites in general (Table 3). Primarily, leeward sites had a markedly higher BSI compared to windward in 2019 but not in 2020 (Table 4). Likewise, bleaching incidence was similar across leeward and windward sites in 2020 (26.96% and 25.63% respectively). Bleaching site susceptibility (BSI) and bleaching incidence was similar across multiple depths and years (56.19% at 6 meters vs. 53.46% at 12 meters in 2019), whereby in 2020 bleaching incidence was higher at greater depth (35.51% against 24.25%). Of the most abundant taxa (n≥10; 13 taxa), seven taxa showed a curvilinear decline in bleaching susceptibility with depth: *Acropora, Favia, Fungia, Galaxea, Goniastrea, Heliopora* and *Leptastrea*, and six taxa: *Pavona, Platygyra, Porites* sp. 1, *Porites* sp. 2, *Psammocora* and *Symphyllia* followed a reversed pattern, bleaching more at intermediate depth (Figure 5). This pattern was consistent across years, except for *Acropora* and *Platygyra*, who showed markedly contrasting bleaching patterns at these depths in 2020. Significant differences in bleaching response across depths were found for *Acropora, Favia, Fungia, Pavona*, and *Porites* sp. 1 in 2019 (Table 5), and for *Acropora, Heliopora, Pavona, Platygyra, Porites* sp. 1 and *Porites* sp. 2 in 2020 (Table 5). Notably, all *Heliopora* colonies bleached at shallow sites in 2019 (Figure 5).

**Table 3.** Bleaching resistance index (BRI) and generalized linear model analysis of selected scleractinian taxa (n≥10) (*note *Echinopora* at leeward sites in 2020 n<10) at leeward and windward sites during successive bleaching events between 2019 and 2020. ***Pr <0.001, **Pr < 0.01, *Pr < 0.05, ns – not significant.

| Genus | leeward 2019 |  | windward 2019 |  | leeward 2020 |  | windward 2020 |  |
| --- | --- | --- | --- | --- | --- | --- | --- | --- |
|  | n | BRI | n | BRI | n | BRI | n | BRI |
| *Acropora* | 164 | 0.76 | 56 | 0.93 | 130 | 0.98 | 41 | 0.98 |
| *Echinopora* * | 19 | 0.14 | 15 | 0.72 | 7* | 0.71 | 17 | 1.00 |
| *Favia* | 11 | 0.87 | 60 | 0.81 | 10 | 1.00 | 104 | 0.95 |
| *Fungia* | 95 | 0.56 | 20 | 0.82 | 107 | 0.90 | 30 | 0.96 |
| *Pavona* | 121 | 0.32 | 23 | 0.91 | 111 | 0.68 | 19 | 1.00 |
| *Porites* sp. 1 | 27 | 0.45 | 175 | 0.48 | 25 | 0.72 | 237 | 0.83 |
| *Porites* sp. 2 | 269 | 0.58 | 178 | 0.78 | 231 | 0.87 | 123 | 0.88 |

| Genus | leeward (LW) vs. windward (WW) 2019 |  |  |  | leeward (LW) vs. windward (WW) 2020 |  |  |  |
| --- | --- | --- | --- | --- | --- | --- | --- | --- |
|  | Estimate | SE | Z value | Pr (>\|z\|) | Estimate | SE | Z value | Pr (>\|z\|) |
| *Acropora* | (WW) -3.50 | 0.42 | -8.280 | *** | (WW) -1.20 | 1.06 | -1.131 | ns |
| *Echinopora* | (WW) -2.83 | 0.93 | -3.057 | ** | (WW) -20.28 | 4300.25 | -0.005 | ns |
| *Favia* | (WW) 0.73 | 0.83 | 0.886 | ns | (WW) 15.71 | 1251.05 | 0.013 | ns |
| *Fungia* | (WW) -3.32 | 0.63 | -5.258 | *** | (WW) -1.55 | 0.77 | -2.030 | * |
| *Pavona* | (WW) -3.29 | 0.77 | -4.291 | *** | (WW) -18.97 | 1496.40 | -0.013 | ns |
| *Porites* sp. 1 | (WW) -0.31 | 0.45 | -0.699 | ns | (WW) -0.95 | 0.43 | -2.238 | * |
| *Porites* sp. 2 | (WW) -1.03 | 0.20 | -5.116 | *** | (WW) 0.30 | 0.24 | 1.211 | ns |

**Table 4.** Summary of bleaching incidence (percent bleaching of hard coral colonies), Bleaching Resistance Index (BRI), and the Bleaching site susceptibility Index (BSI) at numerous biophysical scales during two successive bleaching events in 2019 and 2020 in Pulau Lang Tengah, Northeast Peninsular Malaysia. Shallow sites range from 5 to 7 meters, and deep sites from 10 to 14 meters of water depth.

| Biophysical scale (year) | Colonies | Incidence (%) | BRI | Density (n/m<sup>2</sup>) | BSI |
| --- | --- | --- | --- | --- | --- |
| Leeward (2019) | 747 | 70.82 | 57 | 24.90 | 48.21 |
| Windward (2019) | 1135 | 44.85 | 67 | 28.38 | 35.56 |
| Leeward (2020) | 690 | 26.96 | 87 | 23.00 | 12.78 |
| Windward (2020) | 1183 | 25.63 | 87 | 29.58 | 12.23 |
| Shallow (2019) | 1203 | 56.19 | 66 | 30.08 | 37.78 |
| Deep (2019) | 679 | 53.46 | 59 | 22.63 | 37.12 |
| Shallow (2020) | 1138 | 24.25 | 89 | 28.45 | 11.57 |
| Deep (2020) | 735 | 35.51 | 84 | 24.50 | 12.76 |
| Leeward Shallow (2019) | 592 | 72.30 | 61 | 29.60 | 58.25 |
| Leeward Deep (2019) | 155 | 65.16 | 45 | 15.50 | 60.86 |
| Windward Shallow (2019) | 611 | 40.59 | 71 | 30.55 | 37.30 |
| Windward Deep (2019) | 524 | 50.00 | 63 | 26.20 | 40.38 |
| Leeward Shallow (2020) | 546 | 20.15 | 90 | 27.30 | 12.05 |
| Leeward Deep (2020) | 144 | 52.78 | 76 | 14.40 | 25.14 |
| Windward Shallow (2020) | 592 | 23.99 | 90 | 29.60 | 12.35 |
| Windward Deep (2020) | 591 | 27.41 | 87 | 29.55 | 13.47 |

**Table 5.** Bleaching Resistance Index (BRI) and generalized linear model analysis of selected scleractinian taxa (n≥10) across multiple depths (shallow 5-7 m; deep 10-14 m) during successive bleaching episodes in June 2019 and June 2020. ***Pr <0.001, **Pr < 0.01, *Pr < 0.05, ns – not significant.

| Genus | *Shallow 2019* |  | *Deep 2019* |  | *Shallow 2020* |  | *Deep 2020* |  |
| --- | --- | --- | --- | --- | --- | --- | --- | --- |
|  | n | BRI | n | BRI | n | BRI | n | BRI |
| *Acropora* | 203 | 80 | 17 | 85 | 156 | 99 | 15 | 88 |
| *Favia* | 28 | 64 | 43 | 94 | 61 | 95 | 53 | 97 |
| *Fungia* | 81 | 59 | 34 | 66 | 89 | 92 | 48 | 90 |
| *Galaxea* | 61 | 97 | 35 | 100 | 67 | 94 | 31 | 97 |
| *Goniastrea* | 36 | 27 | 16 | 50 | 27 | 81 | 17 | 91 |
| *Heliopora* | 58 | 0 | 64 | 34 | 31 | 22 | 57 | 75 |
| *Leptastrea* | 35 | 90 | 34 | 97 | 35 | 88 | 45 | 94 |
| *Pavona* | 71 | 64 | 73 | 17 | 63 | 81 | 67 | 65 |
| *Platygyra* | 35 | 90 | 35 | 75 | 56 | 77 | 47 | 86 |
| *Porites* sp. 1 | 103 | 60 | 99 | 34 | 123 | 90 | 139 | 74 |
| *Porites* sp. 2 | 317 | 68 | 130 | 61 | 247 | 89 | 107 | 86 |
| *Psammocora* | 13 | 98 | 9 | 93 | 20 | 100 | 28 | 100 |
| *Symphyllia* | 12 | 95 | 14 | 93 | 20 | 96 | 14 | 94 |

| Genus | *Shallow (S) vs. Deep (D) 2019* |  |  |  | *Shallow (S) vs. Deep (D) 2020* |  |  |  |
| --- | --- | --- | --- | --- | --- | --- | --- | --- |
|  | Estimate | SE | Z value | Pr(>\|z\|) | Estimate | SE | Z value | Pr(>\|z\|) |
| *Acropora* | (S) 1.60 | 0.53 | 3.003 | ** | (S) -3.00 | 0.70 | -4.314 | *** |
| *Favia* | (S) 2.71 | 0.65 | 4.160 | *** | (S) 0.88 | 0.62 | 1.403 | ns |
| *Fungia* | (S) 1.47 | 0.52 | 2.820 | ** | (S) -0.71 | 0.43 | -1.666 | ns |
| *Galaxea* | (S) 17.68 | 1817.76 | 0.010 | ns | (S) 0.36 | 0.85 | 0.419 | ns |
| *Goniastrea* | (S) 1.00 | 0.64 | 1.555 | ns | (S) 0.80 | 0.69 | 1.160 | ns |
| *Heliopora* | (S) 18.78 | 1412.07 | 0.013 | ns | (S) 3.09 | 0.67 | 4.591 | *** |
| *Leptastrea* | (S) 1.71 | 1.12 | 1.517 | ns | (S) 0.75 | 0.69 | 1.090 | ns |
| *Pavona* | (S) -1.75 | 0.38 | -4.581 | *** | (S) -1.77 | 0.39 | -4.579 | *** |
| *Platygyra* | (S) 1.07 | 0.60 | 1.781 | ns | (S) 0.98 | 0.41 | 2.394 | * |
| *Porites* sp. 1 | (S) -1.71 | 0.33 | -5.154 | *** | (S) -1.22 | 0.28 | -4.364 | *** |
| *Porites* sp. 2 | (S) -0.29 | 0.21 | -1.373 | ns | (S) -0.66 | 0.25 | -2.663 | ** |
| *Psammocora* | (S) -0.33 | 1.21 | -0.270 | ns | -7.351e-15 | 1.043e+05 | 0.00 | ns |
| *Symphyllia* | (S) -0.51 | 0.87 | -0.588 | ns | (S) -0.41 | 1.07 | -0.380 | ns |

**Figure 5.**
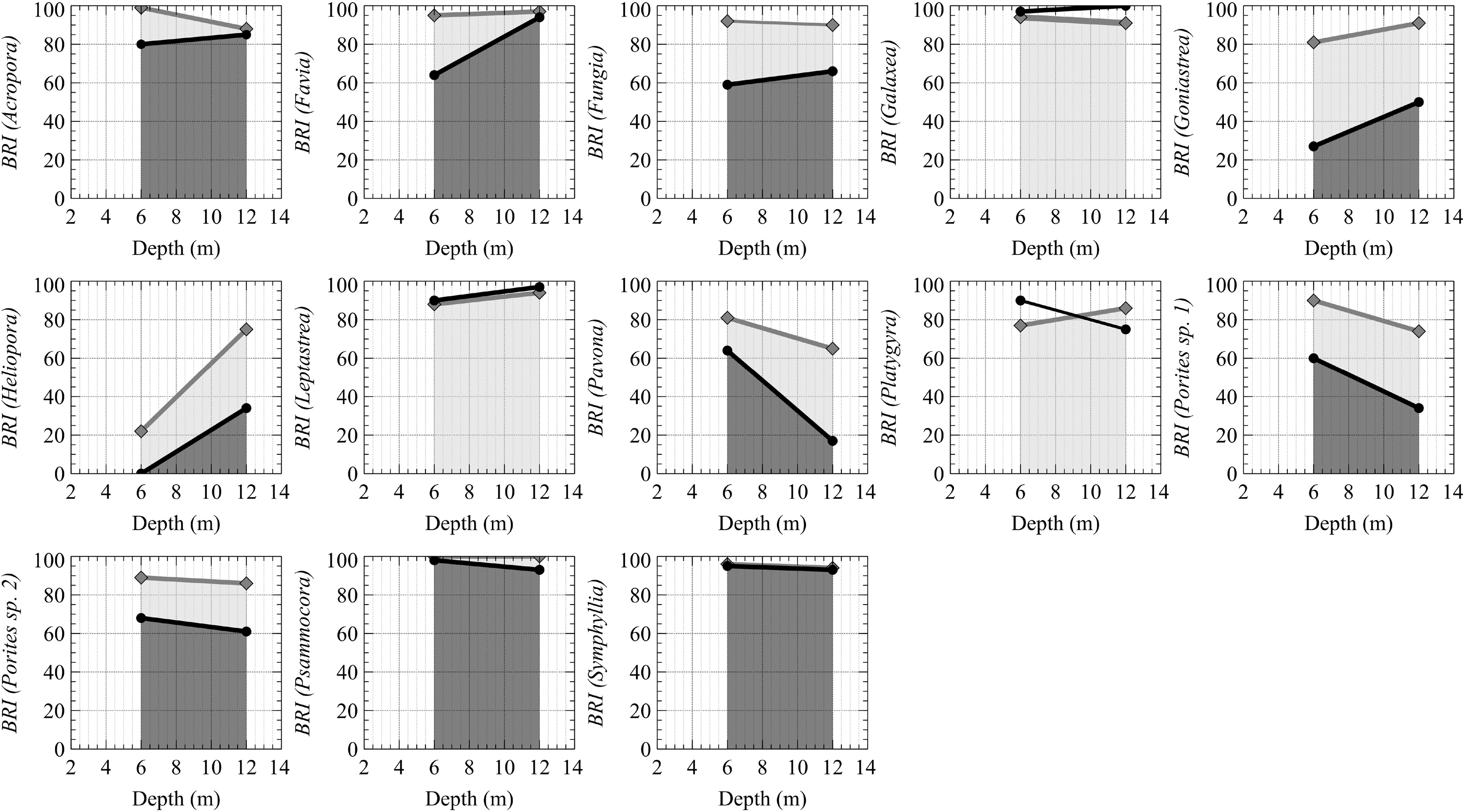
Bleaching Resistance Index (BRI) of scleractinian taxa as a function of depth. Taxa with at least ten observations at each depth are shown. The black lines present the BRI in 2019, and grey lines show the BRI in 2020.

On genus level, Tukey’s post-hoc analysis revealed significant differences between wind and depth interactions for numerous taxa (Table 6). Bleaching response was tested for five taxa with n≥10 (*Acropora, Fungia, Pavona, Porites* sp. 1 and *Porites* sp. 2), and all five taxa were substantially less bleached in June 2019 at shallow windward sites as compared to other sites (Table 6). Apart from *Pavona*, this interaction was significant for all taxa in 2019, especially when compared to shallow leeward sites (Table 6). In particular, *Porites* sp. 2, showed markedly less bleaching in both years at shallow windward locations. Furthermore, *Porites* sp. 1 was significantly different from shallow leeward sites as well deeper windward sites in 2019, and significantly different from deeper windward sites in 2020. Despite significant difference across depth in general for *Porites* sp. 1 (bleaching more at intermediate depth), there was no significant difference between windward deep and leeward shallow in either year for *Porites* sp. 1 and *Porites* sp. 2, highlighting the interactive effect of depth and wind on bleaching susceptibility. Similarly, significant differences across depth were found for *Pavona*, bleaching more at greater depth (Table 5 and Figure 5), whereas differences in bleaching susceptibility of *Pavona* at leeward deep (BRI=10) was chiefly more pronounced compared to windward deep (BRI=80) (Table 6). At deeper leeward sites, *Pavona* showed a very severe bleaching response, and was significantly more bleached here in both years as compared to shallow leeward sites.

**Table 6.**
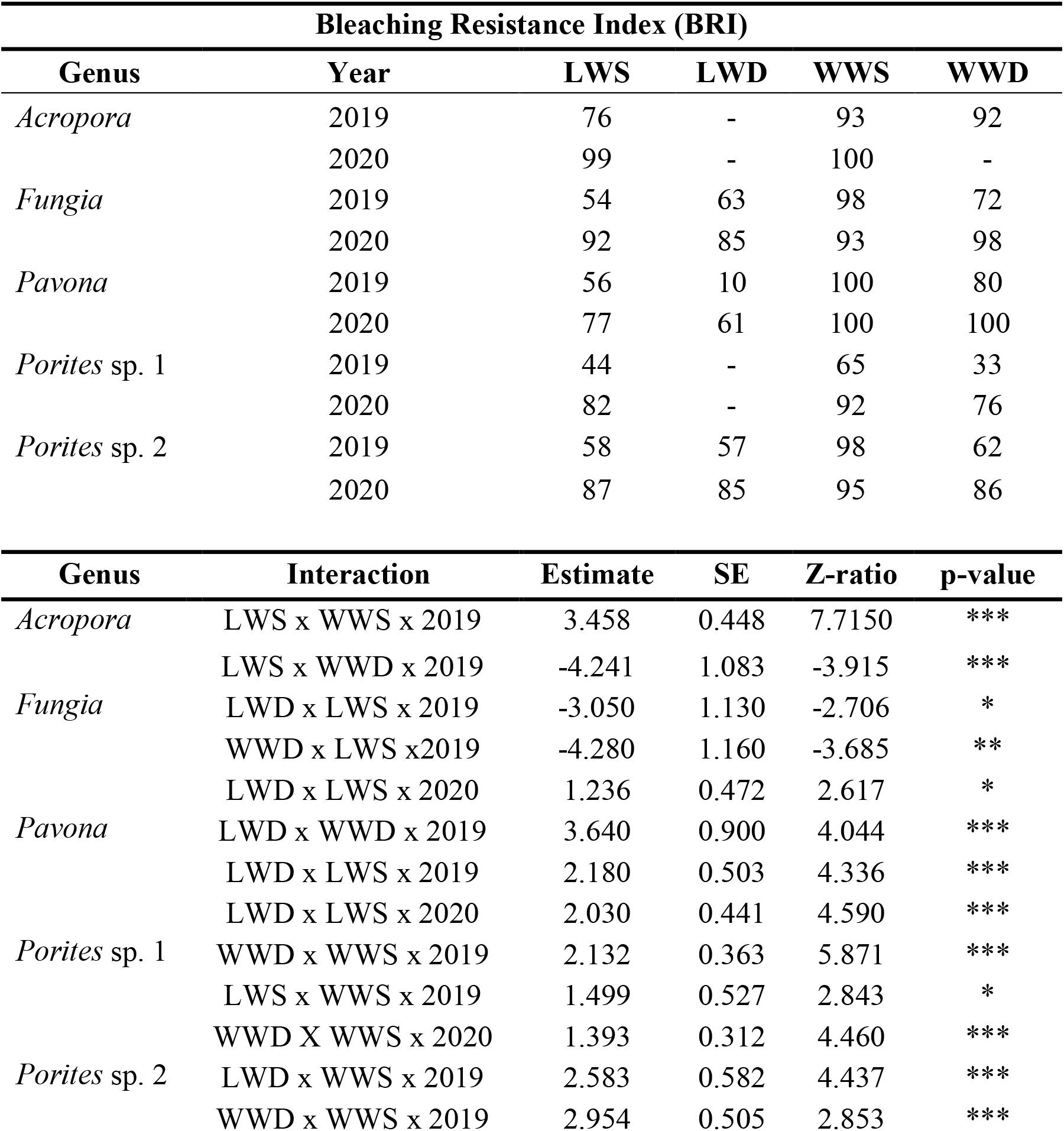
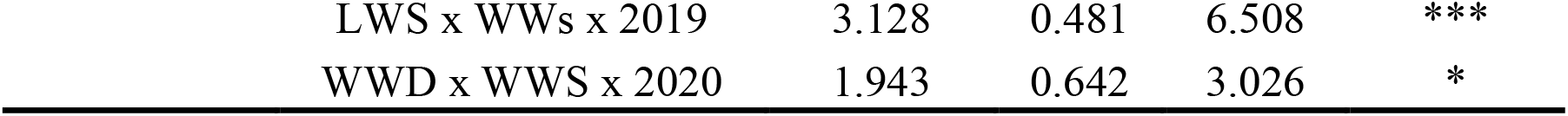
Bleaching resistance Index (BRI) of selected taxa (n≥10) at fine biophysical reef scale (LWS-leeward shallow; LWD-leeward deep; WWS-windward shallow; WWD-windward deep) are shown for two successive bleaching episodes in 2019 and 2020. Tukey’s post-hoc test results are shown for significantly different interactions at significance levels of ***p<0.001, **p<0.01 and *p<0.05 (missing entries are due to insufficient sample size).

## Discussion

### Historical sea surface temperature analysis

Linear regression analysis revealed a warming trend since 1985 (Figure 2) of 0.17°C per decade, while this warming trend is more pronounced during the bleaching seasons (0.20°C). Since 1986, the ocean has absorbed up to eight times as much energy-derived heat as previously between 1958 and 1986, resulting in uninterrupted warming (Cheng et al., 2021). Since industrialization, the oceans have warmed by 1°C until the early 2000s (Stocker et al., 2014; Pachauri et al., 2014, (IPCC)). Overall, this analysis found an expeditious and pertinent warming of 0.58°C since 1985. Analogous to globally accelerating warming rates over the past decade (Skirving et al., 2019), warming rates accelerated locally over the last decade with nine of the ten warmest years being recorded since 2010 (Figure 2; Supplementary S2), whereby 2020 was the second warmest bleaching season on average locally, coinciding with the warmest year of mean ocean temperatures globally (Cheng et al., 2021). Bleaching in 2020 occurred outside of an El Niño Southern-Oscillation (ENSO) associated temperature extreme, furthermore underscoring the diminishing relationship between mass bleaching events and extreme temperature anomalies in ENSO years, chiefly highlighting the impacts of globally observed warming patterns on local reef scale (Hughes et al., 2018b). Based on bleaching susceptibilities of taxa reported here and temperature trends at local and global scale, we argue that mild to moderate coral bleaching (particularly of susceptible taxa) is now likely to occur annually in Northeast Peninsular Malaysia during peak seasonal temperatures between April-September (Figure 2 and 3) and irrelevant of ENSO temperature extremes.

Of 11 thermal anomalies with DHW > 1°C-weeks, six anomalies were recorded over the past decade (Table 1). Bleaching occurrence in Northeast Peninsular Malaysia was reported *in-situ* in 1998 (Kushairi, 1998), in 2010 (Tan and Heron, 2011), in 2014 (Reef Check Malaysia, 2014) and during this study in 2019 and 2020. Furthermore, coral bleaching was observed at mild to moderate levels in Southeast Peninsular Malaysia in 2019 and 2020 (Reef Check Malaysia, 2019; Reef Check Malaysia, 2020) at higher levels of thermal stress (Supplementary S3). In 2016, thermal stress reached a maximum of 1.01°C-weeks in mid-July, but bleaching was not observed *in-situ* in Northeast Peninsular Malaysia (Szereday, personal observation) and neither in Southeast Peninsular Malaysia (Affendi, personal observation). In 1998, coral bleaching in Malaysia was reported for the first time reaching moderate levels (≤30% bleaching incidence), and resulting in low level mortality (Kushairi, 1998). Coral bleaching was severest in 2010, with coral bleaching incidence >50%, and severe full colony bleaching was observed for >66% of coral colonies at DHW levels of above 5°C-weeks (Tan and Heron, 2011). Moreover, coral bleaching in 2010 was observed until 25 m of water depth, and mortality levels were significant (Guest et al., 2012). *In-situ* field observations of coral bleaching occurrence in 1987, 1990, 1991, 2001 and 2013, when thermal anomalies exceeded >1°C-weeks, are not available and it is unknown whether bleaching occurred in these years. Observations from 2016 further suggest that bleaching does not occur universally at DHW > >1°C-weeks, as confounding environmental factors (e.g., irradiance, water flow, vertical mixing and nutrient availability) are likely co-determining thermal stress experienced *in-situ* (Brown, 1997). The Southwest Monsoon starts annually in June and results in strong winds, storms and upwelling (Akhir et al., 2015), and likely plays a factor in cooling seawater temperatures after the annual SST peak in May and June. This may have prevented coral bleaching in 2016 as thermal stress (resulting from extreme ENSO) started to rise during the onset of the Southwest Monsoon, rather than in April and May as in 2010 and 2019, when mass bleaching was observed. Nonetheless, there is reported evidence suggesting that numerous bleaching events and associated mortality have occurred in recent history (1998, 2010 and 2014), possibly influencing thermal tolerance in surviving colonies (*sensu* legacy effect). The 2014-2016 ENSO events resulted in the most severe and widespread pan-tropical coral bleaching event (Eakin et al., 2019), whereby in 2014 some degree of bleaching induced mortality is plausible (Reef Check Malaysia, 2014). These field observations and studies suggest that DHW >1°C-weeks likely represent a gross underestimation of thermal stress experienced *in-situ* by hard corals in Northeast Peninsular Malaysia, but are locally a useful indicator to highlights periods of positive thermal anomalies that are commonly induced by global weather extremes such as ENSO. Ultimately, mean seawater temperatures are perpetually rising around Pulau Lang Tengah (Figure 2), in addition to higher frequencies of thermal anomalies that surpass the thermal maxima of scleractinian taxa, causing repeated thermal stress. Regardless of differences in thermal tolerance and low mortality rates post bleaching events as those reported here (Figure 4 and Supplementary S5), such warming trends and resulting bleaching events selectively pressure scleractinian communities by altering scleractinian community assemblages (Loya et al., 2001; Baker et al., 2008, Hughes et al., 2018a), and likely influence demographic processes in surviving colonies who suffer high bleaching severity, such as massive *Porites* (Morais et al., 2021). Thus, severe annual coral bleaching will represent an imminent risk to coral reef survival and integrity (Van Hooidonk et al., 2017 (UNEP)). Immediate action to reduce greenhouse gas emissions is the only effective long-term conservation strategy in order to prevent systematic coral reef collapse (Van Hooidonk et al., 2016; Frieler et al., 2013; Hughes et al., 2017).

### Taxon-specific variability in annual bleaching response and recovery

The successive bleaching events of 2019 and 2020 represent the climax of a decade of rapid ocean warming (Table 1 and Figure 2). There is no indication that recent and more frequent exposure to heat stress above physiological thresholds and subsequent bleaching (i.e., 2010, 2013, 2014 and 2016) resulted in widespread thermal resilience, as compared to previous observations of local mass bleaching in 2010 (Tan and Heron, 2011). Contrarily, this study reveals taxon-specific variations in thermal susceptibility similar to those in Southeast Peninsular Malaysia and Singapore during the 2010 mass bleaching event (Guest et al., 2012; Guest et al., 2016). Bleaching incidence and intensity was higher in 2019 compared to 2010 for numerous major taxa despite less heat stress in 2019 (Supplementary S6). For instance, eight taxa (*Cyphastrea, Echinopora, Fungia, Favites, Goniastrea, Pocillopora, Porites* sp. 1 and *Porites* sp. 2) showed higher bleaching response at lower levels of heat stress in 2019, whereas the bleaching response of *Acropora, Favia, Galaxea, Montipora, Pavona, Platygyra*, and *Symphyllia* was similar to 2010. In addition, heat stress driven mortality rates were higher only for four taxa (*Acropora, Pocillopora, Pavona, Montipora)* in 2010 compared to 2019, which is readily explained by the DHW theory, where mortality rates increase with higher DHW. In general, mortality rates observed during this study did not substantially differ from expected levels of mortality due to other natural causes (e.g., predation, disease, physical breakage, etc.) (Harriot, 1985). However, ecological footprints of heat disturbance events are not equal due to varying magnitude, severity and spatial extent of thermal events, with biophysical factors further influencing bleaching response at reef scales, possibly underlining the differences observed between Northeast and Southeast Peninsular Malaysia in 2010 and 2019. Nevertheless, we report high bleaching incidence and intensity for taxa such as *Porites* sp. 1, *Porites* sp. 2, *Goniastrea* and *Cyphastrea*, and higher bleaching response at lower levels of heat stress may point to worrying trends where certain scleractinian taxa increase their sensitivity after frequent events of heat disturbance. Of all taxa with n≥10, *Psammocora, Acropora, Galaxea, Leptastrea* and *Sympyhllia* were the most thermally resistant taxa across both years, whereas *Psammocora* was the only taxa to not bleach in 2020. Despite lower heat stress in 2020, *Galaxea, Leptastrea, Platygyra* (in shallow reef sections) and Fungia (at shallow windward reefs) were moderately more susceptible to bleaching in 2020, while the bleaching response of *Acropora* (in deeper reef sections) and *Symphyllia* remained unchanged, suggesting that annual sequences will selectively pressure community assemblages and reduce thermal tolerance in certain taxa. Experiments on species-specific trends have demonstrated that annual exposure can weaken cellular processes involved in thermal tolerance (Grottoli et al., 2014; Schoepf et al., 2015), turning “winners” into “losers” or vice versa (Van Woesik et al., 2011). Our findings suggest that while fast-growing taxa such as *Acropora* and *Montipora* may adapt to higher thermal stress by means of directional selection and reproduction (Figure 4), other taxa possibly do not possess similar life-history strategies or quick adaptive responses due to physiological differences involved in phenotypic plasticity (Darling et al. 2012; Thomas et al., 2018). Such species-specific differences might explain the diverging responses of taxa seen in this study. For instance, two taxa (*Galaxea*, and *Leptastrea*) were moderately more susceptible to bleaching in year 2020, whereas *Cyphastrea, Echinopora*, and *Goniastrea* were more bleaching tolerant. *Porites* sp. 1, *Pavona, Fungia*, and *Favia* bleached considerably less in 2020 (Table 2), albeit accumulated heat stress was lower in 2020 (Figure 3A). *Pocillopora* represented the only major reef-building taxa with fast growth and reproduction, not showing higher bleaching tolerance as historically known (Pratchett et al., 2013; Heron et al., 2016a). As high bleaching incidence and susceptibility are reasoned with levels of heat stress experienced (Liu et al., 2003; Kayanne, 2017), it can be considered that the stronger bleaching response of *Galaxea* and *Leptastrea*, observed in 2020 would further increase to significantly different levels when exposed to heat stress similar to 2019. Likewise, bleaching levels of taxa with lower bleaching susceptibility in 2020 may increase under thermal stress levels as observed in 2019. For example, when conferring *in-situ* models for 2020, differences in thermal stress across both episodes were discernable (1.05°C-weeks vs 0.18°C-weeks at the point of surveying), and differences in heating rates and accumulation prior to bleaching were evident for the 60 day period prior to bleaching (2019 DHD_60_=41.33, HR_60_=0.69 vs. 2020 DHD_60_=27.35, HR_60_=0.46). Thus, heat stress onset was more expeditious and this may have magnified bleaching severity in 2019 (Fordyce et al., 2019). Nonetheless, we cannot suggest that the first bleaching event resulted in any significant strengthening interaction (*sensu* positive legacy), as heat stress levels during the second event did not equal those in 2020, according to satellite measurements. Rather, hard corals continued to be vulnerable to high average temperatures and minor deviations above their thermal threshold, whereas the exact threshold values remain unknown. For taxa such as *Galaxea* and *Leptastrea*, however, a negative legacy effect may have resulted in moderately higher bleaching levels in 2020 under lower levels of heat stress.

In contrast to the above, two taxa (*Acropora*, and *Montipora*) displayed contrasting bleaching hierarchies and improved thermal tolerance as compared historical records (McClanahan et al., 2004; Guest et al., 2012; Pratchett et al., 2013), suggesting thermal adaptation due to directional selection, rather than thermal acclimation. Whereas species-specific genetic and cellular process are underpinning heat stress response in hard corals (Grottoli et al., 2014; Schoepf et al., 2015; Guzman et al., 2018), there is strong evidence that certain functional traits required for establishing thermally tolerant populations after large scale disturbance events are phylogenetically conserved on genus level (Darling et al., 2012; Morais et al., 2021). Such traits include growth, fecundity and reproductive mode, and further supporting evidence is presented by Alvarez-Noriega et al. (2016), who suggest that fecundity can be predicted by coral colony morphology. In this study, we observed scleractinian taxa that possess these functional prerequisites (such as *Acropora*, and *Montipora*), to have reversed their historical bleaching hierarchy, based on low rates of bleaching severity and low rates of bleaching associated mortality. Supposedly, preceding bleaching events eliminated thermally susceptible individuals from the population. Subsequently, successful reproduction and recruitment of the surviving genotypes and taxa with fast growth and reproduction (Darling et al., 2012), perchance resulted in a directional selection since 1998 and 2010 of thermally tolerant genotypes of taxa historically classified as thermally susceptible (Edmunds, 1994; Thompson and van Woesik, 2009). Dramatic short-term recovery (e.g., <5 years) of depleted *Acropora* populations after bleaching events in 2016 and 2017 has been evidently highlighted on the Great Barrier Reef, whereby genus-wide recovery was facilitated by traits such as quick growth and reproduction (Morais et al., 2021). Furthermore, recent studies suggest that thermal stress does not impede heritability of functional traits responsible for growth and survival (Bairos-Novak et al., 2021), and greater thermal plasticity in surviving offspring has been demonstrated (Wong et al., 2021). Nonetheless, populations of *Acropora* and *Montipora* (as well as other historically susceptible taxa) have likely severely declined compared to pre-2010 levels across the Malaysian Peninsular (Toda et al., 2007), leaving behind large fields of coral rubble unsuitable for coral recruitment (Szereday, unpublished data). Thus, thermal adaptation of these taxa as a result of directional selection is possibly offset by unfavorable conditions (e.g., substrate availability and monsoon waves) for substrate recolonization and surviving populations will regionally struggle to recover to pre-disturbance levels.

One notable finding of this study is the high bleaching susceptibility of *Heliopora spp*. (Table 2). In both years, *Heliopora* showed the highest bleaching incidence, severity and mortality of all surveyed taxa, in particular at shallow sites (Table 5, Figure 5 and Supplementary S7). A wide range of field studies conducted in numerous Indo- and Central-Pacific regions (e.g., Palau, Japan, Western Australia, Kiribati) as well as the Indian Ocean (e.g., the Andaman Sea and the Maldives), repeatedly classified this taxon as heat tolerant (Paulay and Benayahu, 1999; Kayanne et al., 2002; Schumacher et al., 2005; Donner et al., 2010; Phongsuwan and Changsang, 2012; Harri et al., 2014; Richards et al., 2018). Supposedly, accretion rates of *Heliopora* increase during positive thermal anomalies, resulting in competitive advantages over scleractinian taxa (Atrigenio et al., 2017; Guzman et al., 2019). Therefore, it has been suggested that species of *Heliopora* may replace scleractinian taxa in the future as dominant reef-builders (Richards et al., 2018). To our knowledge, our findings are the first to report high bleaching susceptibility of *Heliopora sp*., and further research is warranted to elucidate mechanisms that may have reduced its thermal tolerance in Northeast Peninsular Malaysia. Firstly, environmental factors such as irradiance may play a factor. Bleaching levels of *Heliopora* were severe at both depths surveyed. However, bleaching severity was significantly less in deeper reefs in 2019 and 2020 (Table 5 and Figure 5). This is paradoxical, as *Heliopora* is well-known to inhabit shallow reef areas in living and fossil assemblages (Colgan, 1984; Zann and Bolton, 1985). Secondly, symbiotic partnerships may be a driving factors. In the Andaman Sea, *Heliopora* has thus far been identified to host the thermally tolerant zooxanthellae clade D2 (Lajeunesse et al., 2010), whereas generalist clades of *Symbiodinium sp*. have been identified in association with both *Heliopora* species in Western Australia (Clade C1), and clade C3 is known from the central Pacific and the Philippines (Pochon et al., 2010; Guzman et al., 2018; Richards et al., 2018). Moreover, Guzman et al. (2018) identified an array of genes (e.g., heat shock protein and antioxidants) in *Heliopora coerulea* that are involved in environmental stress response, and mixotrophy is widespread in octocorals to reduce reliance on symbiotic partnerships (Fabricius and Klumpp, 1995). Further research is necessary to understand whether the observed susceptibility of *Heliopora* in Pulau Lang Tengah is associated to any of the above factors, and whether *Heliopora* is generally a bleaching susceptible taxon in Northeast Peninsular Malaysia and the Sunda Shelf ecoregion.

Perchance, high warming rates and more frequent thermal stress impairs thermal tolerance of certain taxa, such as *Heliopora*, which cannot meet the required rates for thermal acclimation (Donner et al., 2005). Donner at al., (2005) suggested that in order to survive and acclimate to warming trajectories, hard corals must acclimate at a rate of 0.2-1.0°C per decade, whereas this rate likely increases as ocean warming and return-frequencies of heat disturbance intensify. In addition, Guest et al. (2012) argue that faster thermal acclimation rates of scleractinian taxa in southern Peninsular Malaysia compared to reefs in northern Sumatra were arguably co-facilitated by lower warming rates (0.009°C vs. 0.0161°C). Additionally, higher warming rates are increasing the return frequency of high-temperature anomalies resulting in more frequent heat disturbances, and are simultaneously narrowing DTR, as the temperature minimum increases (Figure 2A). Eventually, hard corals in Northeast Peninsular Malaysia are exposed to their upper physiological thermal threshold longer and more frequently, possibly impacting other physiological and cellular processes and functions to the detriment of general colony fitness. In summary, bleaching responses of scleractinian taxa vary among time and space (McClanahan 2004), whereby bleaching susceptibility and thermal tolerance are not equivocal across larger spatial scales and may be reversed when primary stressors agents such as frequent heat stress become overly prevalent. Ultimately, annual coral bleaching results in selective pressure even when thermal stress is not sufficient to trigger mass mortality (Hughes et al., 2019), and can counteract acclimation potential of taxa based on species-specific traits (Grottoli et al., 2014; Schoepf et al., 2015), and finally ameliorates diverging bleaching responses if heat stress will continue to increase in magnitude and frequency.

### Bleaching response on biophysical reef scale

*In-situ* measurements revealed significant differences in diel temperature profiles across windward and leeward reef sites (Figure 3, Supplementary 4), and generalized linear model (GLM) bleaching analysis of taxa revealed significant difference across depth and wind exposure (Table 3 and 5), whereby the combined interactions of wind and depth (e.g., leeward shallow) revealed complex abiotic-biophysical networking (Table 6), resulting in lower bleaching response at shallow windward sites. This contrasts previous research that reported leeward coral reef sites are more thermally tolerant (McClanahan and Maina, 2003; Voolstra et al., 2020). Thermal regimes defined as high-frequency diel temperature variations (DTR), were significantly more variable at leeward sites for all seasonal time series (Supplementary S4), apart from the cooler monsoon season (DTR_FW_ Kruskal-Wallis H=1.17, p=0.56). Greater DTR ranges at leeward sites are commonly documented (Safie et al., 2018), and were suggested to benefit thermal tolerance in hard corals at reef scale (Oliver and Palumbi, 2011; Thomas et al., 2018). The observed variations in DTR across sites were largely driven by higher average maximum temperatures at leeward sites (Figure 3, Supplementary S4). In example, during the bleaching season, average maximum temperatures were 31.06, 30.93 and 31.00 at HR, BK1 and TT1, respectively. These differences were more pronounced during the peak summer season and during the 60 days period prior to bleaching. Therefore, greater bleaching response of taxa at leeward sites and higher site-specific susceptibility regardless of depth (Table 4), is likely due to greater overall heat accumulation at leeward sites as indicated by higher average daily maximum temperature. Whereas some of these differences in bleaching response can at least partially be attributed to species identity (see below), wind exposure (Figure 1B) and subsequent water mixing and reduced irradiance are likely reducing heat accumulation during the peak bleaching season at windward sites. This is further supported by the fact that differences in DTR across sites were not statistically significant during the more turbulent monsoon season (DTR_FW_) (when powerful monsoon winds impact all sites more equally), thus indicating significant seasonal variations that result in greater heat accumulation at leeward sites. These fine differences in scleractinian bleaching response at marginal differences of thermal stress point to a variety of confounding factors that influence and induce coral bleaching. For instance, cloud cover and subsequent irradiance, water flow and vertical mixing may significantly magnify bleaching stress even under low levels of heat stress (Nakamura and van Woesik, 2001; Page et al., 2019), and anthropogenic eutrophication (here HR) likely reduces bleaching resilience in hard corals (Wooldridge et al., 2012). Such factors may have acted synergistically with high seawater temperatures in 2019 and 2020, and this study did not investigate such environmental parameters directly. Rather, we conducted reef scale surveys and bleaching response analysis as to be able to proxy the effects of irradiance and vertical mixing, by recording bleaching response as a function of depth and wind (Table 3 and 5). In further steps, the analysis of depth and wind as interactive effects revealed substantial differences in bleaching response for several taxa across fine biophysical scales (Table 4 and 6). Therefore, we suggest a complex networking effect of wind, light and depth on bleaching response across and within individual reefs, further underlined by high-frequency diel temperature variability (DTR). This complex network of environmental differences may further inlfuence species-specific physiological limits that determine adaptive mechanism and upper thermal limits, such as genotypic predisposition (Dixon et al. 2015), host-specific physiological traits (Fitt et al., 2009), and associations with specific symbionts and holobionts (Baker et al., 2004; Voolstra et al., 2021). Thus, species identity played a factor for numerous, but not all taxa. Taxonomically driven susceptibilities may be steered by environmental forces impacting biological stress expressions, while determining species distribution along environmental gradients. Shifts in species composition can be found with increasing depth (Roberts et al., 2015), and differentiated bleaching susceptibility of species at multiple depths is common (Bridge et al., 2014; Muir et al., 2017; Muñiz-Castillo and Arias-Gonzalez, 2021). Here, two patterns of bleaching response across multiple depths were found: either a constant decline with increasing depth (seven taxa), or an increase of bleaching severity with increasing depth (six taxa). Baird et al. (2018) found that bleaching incidence declined for most taxa with depth on the Great Barrier Reef in 2016. In contrast, studies on oceanic reefs in Australia found several pronounced non-curvilinear responses of taxa similar to our findings (Crosbie et al., 2019). Particularly the bleaching patterns of *Porites* sp. 1 were strikingly similar, bleaching more severely at intermediate depth. Likewise, patterns seen here for *Pavon*a are reflecting patterns seen by Baird et al. (2018), where bleaching response increased at intermediate depth. In these incidences, it is conceivable that single environmental factors present at shallow reef sections, such as light irradiance (Brown et al., 2002), enhance adaptive response (acclimation) to an overarching environmental variable such as diel temperature variability for some taxa. During events such as an extreme ENSO, shallow water acclimation may be beneficial, but the bleaching responses observed *in-situ* are the result of interactions between multiple agents (i.e., wind, light, temperature). Particularly for *Porites* sp.1 and sp. 2, our findings suggest that the interactive effects of wind and shallow water (in proxy of irradiance) are underlining differences observed across finer biophysical gradients at individual reef scales (Table 6), ultimately leading to overall greater thermal plasticity of shallow colonies as compared to colonies found at the same site, but at greater depth (Baird et al., 2018; Crosbie et al., 2019). Whereas diel temperature variability and high light irradiance in shallow water possibly induce acclamatory mechanism to heat stress (Brown et al., 2002), during blanket heat stress periods (such as ENSO extremes), vertical mixing may act as a protective agent by increasing water flow and light scattering (Fordyce et al., 2019; Page et al., 2019). Subsequently, wind exposure plays an additional role in determining bleaching incidence and severity by modulating light attenuation and water flow at reef scale (Lesser, 1997; Nakamura and van Woesik, 2001). Windward exposure significantly impacted the bleaching responses of *Acropora, Echinopora, Pavona, Porites* sp. 1 and *Porites* sp. 2 (Table 3), bleaching significantly less at windward sites.

The interactive effects of depth and wind on bleaching response were tested for taxa with sufficient observations and compared interactions individually (Table 6). At fine biophysical reef scale, numerous taxa exhibited significant differences in bleaching susceptibility. Notably, *Pavona* was particularly tolerant at shallow windward sites and especially susceptible at deeper leeward sites (Table 6). The latter is chiefly exposed to sedimentation stress, which possibly magnified the bleaching response observed. Additionally, species-specific thermal response may have determined high site-specific susceptibility, as surveyed *Pavona* at both leeward depths were exclusively foliose *Pavona (e*.*g*., *Pavona cactus)*, which dominated the scleractinian assemblage at this scale in both years (n 2019 = 59 of 144, n 2020 = 63 of 155). At windward sites, however, *Pavona* was mainly only found in encrusting forms, and thus the observed differences in bleaching response for this taxon is likely a result of taxon-identity and sedimentation. Moreover, *Acropora* was more susceptible at shallow leeward sites and mostly bleached at mild levels (e.g., fluorescing), whereby this likely represents a species-specific susceptibility as most *Acropora* recorded here are hispidose *Acropora* (e.g., *Acropora longicyathus*, see Supplememtary S5). This morpho-taxon was almost exclusively recorded at shallow leeward sites (140 of 145 colonies recorded). Apart from foliose *Pavona* and hispidose *Acropora*, further significant taxon-specific variations in bleaching susceptibility were found at biophysical scales (Table 4), which are likely not explained by species identity alone. For instance, abundant taxa such as *Fungia, Porites* sp. 1 and *Porites* sp. 2 were significantly more susceptible to bleaching at shallow leeward than at shallow windward sites. Particularly, *Porites* sp. 1 and *Porites* sp. 2 showed significant differences across windward and leeward sites, across depth in general, and across depth within the same reef (Table 3, Table 5 and Figure 5). Therefore, bleaching response was consistently lowest at shallow windward sites (Table 6), as the combined effects of wind and depth are possibly overarching the isolated effects of wind and depth on bleaching response of these taxa. For example, comparative Tukey’s post-hoc tests of windward deep and windward shallow, as well as leeward shallow and windward shallow were significant for *Porites* sp. 1, whereas leeward shallow and windward deep were not significantly different, indicating similar thermal thresholds of *Porites* sp. 1 at leeward shallow and windward deep, despite clear bleaching patterns across depths. This highlights higher bleaching thresholds at shallow windward sites that over mask bleaching patterns across depth, emphasizing variability to thermal stress within individual sites. Similarly, *Porites* sp. 2 showed significantly higher bleaching tolerance at shallow windward sites during both bleaching events (Table 6). Although only these five taxa were tested due to sample size, these results strongly suggest that at least some taxa are more prone to coral bleaching at shallow sites with less wind exposure. The observed differences for the two *Porites* groups clearly underscore an ample spectrum of taxon-specific bleaching susceptibility on the scale of individual reefs.

In conclusion, high-frequency diel temperature variability does not universally result in higher thermal tolerance during extreme events (*sensu* acclimation to a single variable). Rather, reef-specific wind exposure, in interplay with diel temperature variability, significantly modulated bleaching response due to complex networking of environmental and biological interactions (Sugget and Smith, 2019). Further investigations focusing on species-specific susceptibility across a larger sampling area, in combination with detailed environmental data and temperature measurements across multiple depths are required to better understand the interactive effects of wind, depth, temperature and local site pressures (e.g., sewage pollution) on taxon- and species-specific bleaching susceptibility at biophysical reef scale to potently identify bleaching tipping points and upper physiological temperature limits. We do not present *in-situ* temperature measurements across multiple depths at these sites, but temperature differences across depth are inherent. Additionally, there were notable discrepancies in the distribution of taxa across such wind and depth environments (Table 3; Table 5), resulting in few taxa being examined and skewing results due to sample sizes. Taxa that were equally abundant across wind sites are *Echinopora* and *Porites* sp. 2 (in 2019), and seven taxa were equally distributed across depth: *Favia, Heliopora, Leptastrea, Platygyra, Pavona, Porites* sp. 1 and *Symphyllia*. Lastly, it cannot be ruled out that some of the contrasting responses are not at least partially due to sewage discharge by nearby beach resorts, as such local pressure may magnify bleaching severity by reducing colony fitness in general (Wooldridge et al., 2012). This study does not present records of nutrient content across leeward and windward sites, but impacts from sewage discharge are unlikely at windward sites as these are not in close proximity to river estuaries, coastal development or other similar sources of pollution and sedimentation.

### Bleaching thresholds at reef scale: *in-situ* vs. satellite temperature monitoring

The 2019 and 2020 thermal events were similar in magnitude and exceeded the thermal maxima of scleractinian communities. Remotely sensed and *in-situ* measurements were closely correlated (annual mean Pearson r=0.97, p<0.001; bleaching season Pearson r=0.76, p<0.001). However, the observed average differences between measurement tools during the 2020 bleaching season

(0.63±0.26°C) suggests that remotely sensed measurements are possibly misestimating accumulated heat stress, similar to reports by Guest et al. (2012) in Southeast Peninsular Malaysia. Although satellites measurements are well-suited to determine heat stress, bleaching indices and temperature trends on larger scale (Heron et al., 2014), choosing the best-fit model and most accurate bleaching indices remains a challenge when historical measurements are absent, especially over fine spatial and temporal scales. Degree-heating weeks is an operational web-based product at a global scale. Remotely sensed temperature data is recorded by multiple satellites and sensors and is calculated by an algorithm combining past and present data within and beyond the selected 5 km^2^ grid. Consequently, due to changes in algorithm, sensors, and individual satellite accuracy, remotely sensed measurements represent an estimation of the real temperature experienced at any given location (Heron et al., 2014; Liu et al., 2014). In contrast, *in-situ* measurements, are accurate point measurements. In 2019, 55.21% of all surveyed colonies bleached, with 30.29% of colonies bleaching very severely (>66% of colony surface). Such ecologically significant bleaching responses are known for DHW values exceeding at least 4°C-weeks (Eakin et al., 2010; Liu et al., 2014; Heron et al., 2016a). Bleaching thresholds of scleractinian taxa used in this study are based on web-based products, demonstrated to both, under- and overestimate heat stress experienced *in-situ* (McClanahan et al., 2007b; Kayanne, 2017). Partly, such discrepancies are expected errors of satellites recording near-shore (e.g., reflection from landmasses), requiring measurements to be recorded further out at sea (here 2.5 km from the islands shore), where lateral and vertical movement of water reduces SST, compared to *in-situ* reef measurements in shallower and more stagnant coastal waters. Moreover, selecting a “one size fits all’-model is not feasible when considering the amount of highly localized stressors that may exacerbate coral bleaching by reducing overall community fitness (Brown, 1997; Wooldridge et al., 2012). Deriving the MMM by interpolating *in-situ* measurements on satellite data did offer a closer explanation of the bleaching incidence in 2020 than satellite derived data. This Model suggests that scleractinian taxa possess a markedly higher temperature threshold (MMM 30.51°C + 1°C), but lower bleaching resistance, as 0.18°C-weeks (equivalent to two days of 1°C above the MMM) sufficed to trigger widespread and moderate coral bleaching in June 2020 (BRI=88), affecting 17 of the 18 most abundant taxa and 20 of 33 taxa overall. In Japan, low levels of heat stress (e.g., <3°C-weeks) resulted in ecologically significant and widespread bleaching (Strong et al., 2002), underscoring that global threshold values (such as DHW 4°C-weeks) do not apply universally at reef scale. Ultimately, runaway ocean warming may have additionally propelled a higher bleaching sensitivity, as discussed above, since observed bleaching responses do indicate a general sensitivity to above thresholds temperatures.

It cannot be ruled out that thermal thresholds have not been overestimated rather than underestimated. Adding the mean monthly difference of *in-situ* and remote data to satellite measurements for each respective month (e.g., 0.57°C for May 2020 applied to each satellite measurement in May 2020), while keeping the predetermined MMM threshold of 29.94°C +1°C (compared to 31.51°C), would result in DHW of 9.92°C-weeks in 2020 and severe thermal stress. In this scenario, scleractinian taxa demonstrate mechanisms that increase thermal tolerance after experiencing acute thermal stress (Thompson and van Woesik, 2009; Thomas et al., 2018; Fox et al., 2019). Nonetheless, neither model readily explained the bleaching levels observed, whereas immediate impacts from secondary sewage outflow can be ruled out for at least four sites (all windward). In conclusion, longer-term *in-situ* temperature data over environmental gradients combined with field observations of coral bleaching incidence and severity are needed to accurately identify the current thermal baseline and tolerance of scleractinian communities at reef scale in northeastern Peninsular Malaysia. However, based on these findings, a conservative estimate would suggest that bleaching of susceptible taxa starts to occur in Northeast Peninsular Malaysia when temperatures exceed 31.0°C, whereas more resilient taxa start to experience bleaching at >32.0°C. Ultimately, this study provides a valuable diagnostic for the regional understanding of coral bleaching susceptibility, interactive effects of environmental and biological properties, and highlights the severe future consequences of unchecked global warming for tropical marine ecosystems.

## Author contributions

SS designed and conceived the study, conducted the fieldwork, analyzed the data and wrote the original draft. AYA analyzed the data, supported technical framework development and edited the original draft. All authors edited the final draft and granted final approval to submit for publication.

## Acknowledgments

Research was conducted under permit number Prk.ML.630-7Jld.5 (21) issued by the Department of Fisheries (DoF) Malaysia, and permit number 40/200/193711 (2), issued by the Ministry of Economic Affairs (MEA) Malaysia. We are grateful to Albert Apollo Chan and Dato’ Haji Munir bin Haji Mohd. Nawi from the Department of Fisheries Malaysia, as well as to Muhammad Jawad bin Tajuddin (Ministry of Economic Affairs Malaysia), for supporting our research and facilitating necessary research permits. We are thankful to Summer Bay Resort, Lang Tengah Island, for dedicatedly supporting our research and providing access to their facilities. We would like to thank Joseph A. Henry for providing us with the temperature monitors necessary to conduct this study. We are thankful to Gang Liu, Joseph A. Henry, Natasha Zulaikha, and Chew KL. for giving critical suggestions, input and support that greatly improved the manuscript. Lastly, the writing of this article was supported by a writing residency at Rimbun Dahan.

## Footnotes

1 https://globalwindatlas.info/

2 https://pae-paha.pacioos.hawaii.edu/erddap/griddap/dhw_5km.html

## References

Akhir, M. F., Daryabor, F., Husain, M. L., Tangang, F., & Qiao, F. (2015). Evidence of Upwelling along Peninsular Malaysia during Southwest Monsoon. Open Journal of Marine Science, 05(03), 273–279. https://doi.org/10.4236/ojms.2015.53022

Álvarez-Noriega, M., Baird, A. H., Dornelas, M., Madin, J. S., Cumbo, V. R., & Connolly, S. R. (2016). Fecundity and the demographic strategies of coral morphologies. Ecology, 97(12), 3485–3493. https://doi.org/10.1002/ecy.1588

Ainsworth, T. D., Heron, S. F., Ortiz, J. C., Mumby, P. J., Grech, A., Ogawa, D., et al. (2016). Climate change disables coral bleaching protection on the Great Barrier Reef. Science, 352(6283), 338–342. https://doi.org/10.1126/science.aac7125

Atrigenio, M., Aliño, P., & Conaco, C. (2017). Influence of the blue coral Heliopora coerulea on scleractinian coral larval recruitment. Journal of Marine Biology, 2017. https://doi.org/10.1155/2017/6015143

Baker, A. C., Starger, C. J., McClanahan, T. R., & Glynn, P. W. (2004). Corals’ adaptive response to climate change. Nature, 430(7001), 741–741. https://doi.org/10.1038/430741a

Baker, A. C., Glynn, P. W., & Riegl, B. (2008). Climate change and coral reef bleaching: An ecological assessment of long-term impacts, recovery trends and future outlook. Estuarine, Coastal and Shelf Science, 80(4), 435–471. https://doi.org/10.1016/j.ecss.2008.09.003

Barbosa, A. M., Brown, J., Jiménez-Valverde, A., & Real, R. (2014). ModEvA: Model Evaluation and Analysis R Package, Version 1.3

Baird, A. H., Madin, J. S., Álvarez-Noriega, M., Fontoura, L., Kerry, J. T., Kuo, C. Y., et al. (2018). A decline in bleaching suggests that depth can provide a refuge from global warming in most coral taxa. Marine Ecology Progress Series, 603, 257–264. https://doi.org/10.3354/meps12732

Baird, A. H., & Marshall, P. A. (2002). Mortality, growth and reproduction in scleractinian corals following bleaching on the Great Barrier Reef. Marine Ecology Progress Series, 237, 133–141.

Bairos-Novak, K. R., Hoogenboom, M. O., Oppen, M. J. H., & Connolly, S. R. (2021). Coral adaptation to climate change: Meta-analysis reveals high heritability across multiple traits. Global Change Biology, July, 1–17. https://doi.org/10.1111/gcb.15829.

Bollati, E., D’Angelo, C., Alderdice, R., Pratchett, M., Ziegler, M., & Wiedenmann, J. (2020). Optical Feedback Loop Involving Dinoflagellate Symbiont and Scleractinian Host Drives Colorful Coral Bleaching. Current Biology, 30(13), 2433-2445.e3. https://doi.org/10.1016/j.cub.2020.04.055

Bridge, T. C. L., Hoey, A. S., Campbell, S. J., Muttaqin, E., Rudi, E., Fadli, N., et al. (2013). Depth-dependent mortality of reef corals following a severe bleaching event: implications for thermal refuges and population recovery. F1000Research, 2, 187. https://doi.org/10.12688/f1000research.2-187.v1

Brown, B. E. (1997). Coral bleaching: causes and consequences. Coral reefs, 16(1), S129–S138. https://doi.org/10.1007/s003380050249

Brown, B. E., Dunne, R. P., Goodson, M. S., & Douglas, A. E. (2002). Experience shapes the susceptibility of a reef coral to bleaching. Coral Reefs, 21(2), 119–126. https://doi.org/10.1007/s00338-002-0215-z

Cannon, S. E., Aram, E., Beiateuea, T., Kiareti, A., Peter, M., & Donner, S. D. (2021). Coral reefs in the Gilbert Islands of Kiribati: Resistance, resilience, and recovery after more than a decade of multiple stressors. PLoS ONE, 16(8 August), 1–29. https://doi.org/10.1371/journal.pone.0255304

Cheng, L., Abraham, J., Trenberth, K. E., Fasullo, J., Boyer, T., Locarnini, R., et al. (2021). Upper Ocean Temperatures Hit Record High in 2020. Advances in Atmospheric Sciences. https://doi.org/10.1007/s00376-021-0447-x

Crosbie, A. J., Bridge, T. C. L., Jones, G., & Baird, A. H. (2019). Response of reef corals and fish at Osprey Reef to a thermal anomaly across a 30 m depth gradient. Marine Ecology Progress Series, 622, 93–102. https://doi.org/10.3354/meps13015

Darling, E. S., Alvarez-Filip, L., Oliver, T. A., Mcclanahan, T. R., & Côté, I. M. (2012). Evaluating life-history strategies of reef corals from species traits. Ecology Letters, 15(12), 1378–1386. https://doi.org/10.1111/j.1461-0248.2012.01861.x

Dixon, G. B., Davies, S. W., Aglyamova, G. V., Meyer, E., Bay, L. K., & Matz, M. V. (2015). Genomic determinants of coral heat tolerance across latitudes. Science, 348(6242), 1460–1462.

Donner, S. D., Kirata, T., & Vieux, C. (2010). Recovery From the 2004 Coral Bleaching Event in the Gilbert Islands, Kiribati. Atoll Research Bulletin, 587.

Donner, S. D., Skirving, W. J., Little, C. M., Oppenheimer, M., & Hoegh-Gulberg, O. (2005). Global assessment of coral bleaching and required rates of adaptation under climate change. Global Change Biology, 11(12), 2251–2265. https://doi.org/10.1111/j.1365-2486.2005.01073.x

Eakin, C. M., Morgan, J. A., Heron, S. F., Smith, T. B., Liu, G., Alvarez-Filip, L., Baca, B., Bartels, E., et al. (2010). Caribbean corals in crisis: Record thermal stress, bleaching, and mortality in 2005. PLoS ONE, 5(11), e13969. https://doi.org/10.1371/journal.pone.0013969

Eakin, C. M., Sweatman, H. P. A., & Brainard, R. E. (2019). The 2014–2017 global-scale coral bleaching event: insights and impacts. Coral Reefs, 38(4), 539–545. https://doi.org/10.1007/s00338-019-01844-2

Edmunds, P. J. (1994). Evidence that reef-wide patterns of coral bleaching may be the result of the distribution of bleaching-susceptible clones. Marine Biology, 121(1), 137–142.

Edmunds, P. J., & Gates, R. D. (2008). Acclimatization in tropical reef corals. Marine Ecology Progress Series, 361, 307–310. https://doi.org/10.3354/meps07556

Fabricius, K. E., & Klumpp, D. W. (1995). Widespread mixotrophy in reef-inhabiting soft corals: the influence of depth, and colony expansion and contraction on photosynthesis. Marine Ecology Progress Series, 125(1–3), 195–204. https://doi.org/10.3354/meps125195

Fitt, W. K., Gates, R. D., Hoegh-Guldberg, O., Bythell, J. C., Jatkar, A., Grottoli, A. G., et al. (2009). Response of two species of Indo-Pacific corals, Porites cylindrica and Stylophora pistillata, to short-term thermal stress: The host does matter in determining the tolerance of corals to bleaching. Journal of Experimental Marine Biology and Ecology, 373(2), 102– 110. https://doi.org/10.1016/j.jembe.2009.03.011

Fordyce, A. J., Ainsworth, T. D., Heron, S. F., & Leggat, W. (2019). Marine heatwave hotspots in coral reef environments: Physical drivers, ecophysiological outcomes, and impact upon structural complexity. Frontiers in Marine Science, 6, 498. https://doi.org/10.3389/fmars.2019.00498

Fox, J., & Weisberg, S. (2018). An R companion to applied regression. Sage publications.

Fox, M. D., Carter, A. L., Edwards, C. B., Takeshita, Y., Johnson, M. D., Petrovic, V., et al. (2019). Limited coral mortality following acute thermal stress and widespread bleaching on Palmyra Atoll, central Pacific. Coral Reefs, 38(4), 701–712. https://doi.org/10.1007/s00338-019-01796-7

Frieler, K., Meinshausen, M., Golly, A., Mengel, M., Lebek, K., Donner, S. D., et al. (2013). Limiting global warming to 2C is unlikely to save most coral reefs. Nature Climate Change, 3(2), 165–170. https://doi.org/10.1038/nclimate1674

Gintert, B. E., Manzello, D. P., Enochs, I. C., Kolodziej, G., Carlton, R., Gleason, A. C. R., et al. (2018). Marked annual coral bleaching resilience of an inshore patch reef in the Florida Keys: A nugget of hope, aberrance, or last man standing? Coral Reefs, 37(2), 533–547. https://doi.org/10.1007/s00338-018-1678-x

Global Wind Atlas 3.0. (2020). https://globalwindatlas.info/ [Accessed October 24, 2020]

Glynn, P. W., & D’croz, L. (1990). Experimental evidence for high temperature stress as the cause of El Nino-coincident coral mortality. Coral reefs, 8(4), 181–191.

Grottoli, A. G., Warner, M. E., Levas, S. J., Aschaffenburg, M. D., Schoepf, V., Mcginley, M., et al. (2014). The cumulative impact of annual coral bleaching can turn some coral species winners into losers. Global Change Biology, 20(12), 3823–3833. https://doi.org/10.1111/gcb.12658

Guest, J. R., Low, J., Tun, K., Wilson, B., Ng, C., Raingeard, D., … & Steinberg, P. D. (2016). Coral community response to bleaching on a highly disturbed reef. Scientific Reports, 6(1), 1–10. https://doi.org/10.1038/srep20717

Guest, J. R., Baird, A. H., Maynard, J. A., Muttaqin, E., Edwards, A. J., Campbell, S. J., et al. (2012). Contrasting patterns of coral bleaching susceptibility in 2010 suggest an adaptive response to thermal stress. PLoS ONE, 7(3). https://doi.org/10.1371/journal.pone.0033353

Guzman, C., Atrigenio, M., Shinzato, C., Aliño, P., & Conaco, C. (2019). Warm seawater temperature promotes substrate colonization by the blue coral, Heliopora coerulea. PeerJ, 7, e7785. https://doi.org/10.7717/peerj.7785

Guzman, C., Shinzato, C., Lu, T. M., & Conaco, C. (2018). Transcriptome analysis of the reef-building octocoral, Heliopora coerulea. Scientific Reports, 8(1), 1–11. https://doi.org/10.1038/s41598-018-26718-5

Harii, S., Hongo, C., Ishihara, M., Ide, Y., & Kayanne, H. (2014). Impacts of multiple disturbances on coral communities at Ishigaki Island, Okinawa, Japan, during a 15 year survey. Marine Ecology Progress Series, 509, 171–180. https://doi.org/10.3354/meps10890

Harriott, V. J. (1985). Mortality rates of scleractinian corals before and during a mass bleaching event. Marine ecology progress series, 21(1), 81–88.

Heron, S. F., Johnston, L., Liu, G., Geiger, E. F., Maynard, J. A., De La Cour, J. L., et al. (2016). Validation of reef-scale thermal stress satellite products for coral bleaching monitoring. Remote Sensing, 8(1), 59. https://doi.org/10.3390/rs8010059

Heron, S. F., Liu, G., Eakin, C. M., Skirving, W. J., Muller-karger, F. E., Vega-Rodriguez, M., et al. (2015). Climatology Development for NOAA Coral Reef Watch’s 5-km Product Suite. https://doi.org/10.7289/V59C6VBS

Hoegh-Guldberg, O. (2011). Coral reef ecosystems and anthropogenic climate change. Regional Environmental Change, 11(1), 215–227. https://doi.org/10.1007/s10113-010-0189-2

Hoegh-Guldberg, O. (1999). Climate change, coral bleaching and the future of the world’s coral reefs. Marine and freshwater research, 50(8), 839–866. https://doi.org/10.1071/MF99078

Hughes, T. P., Kerry, J. T., Álvarez-Noriega, M., Álvarez-Romero, J. G., Anderson, K. D., Baird, A. H., et al. (2017). Global warming and recurrent mass bleaching of corals. Nature, 543(7645), 373–377. https://doi.org/10.1038/nature21707

Hughes, T. P., Kerry, J. T., Baird, A. H., Connolly, S. R., Dietzel, A., Eakin, C. M., et al. (2018a). Global warming transforms coral reef assemblages. Nature, 556(7702), 492–496. https://doi.org/10.1038/s41586-018-0041-2

Hughes, T. P., Kerry, J. T., Connolly, S. R., Álvarez-Romero, J. G., Eakin, C. M., Heron, S. F., et al. (2021). Emergent properties in the responses of tropical corals to recurrent climate extremes. Current Biology, 31(23), 5393-5399.e3. https://doi.org/10.1016/j.cub.2021.10.046

Hughes, T. P., Kerry, J. T., Connolly, S. R., Baird, A. H., Eakin, C. M., Heron, S. F., et al. (2019). Ecological memory modifies the cumulative impact of recurrent climate extremes. In Nature Climate Change, 9(1), 40–43. https://doi.org/10.1038/s41558-018-0351-2

Hughes, T. P., Anderson, K. D., Connolly, S. R., Heron, S. F., Kerry, J. T., Lough, J. M., et al. (2018b). Spatial and temporal patterns of mass bleaching of corals in the Anthropocene. Science, 359(6371), 80–83. https://doi.org/10.1126/science.aan8048

Jaap, W. C. (1979). Observations on zooxanthellae expulsion at Middle Sambo Reef, Florida Keys. Bulletin of Marine Science, 29(3), 414–422.

Johnstone, J. F., Allen, C. D., Franklin, J. F., Frelich, L. E., Harvey, B. J., Higuera, P. E., et al. (2016). Changing disturbance regimes, ecological memory, and forest resilience. Frontiers in Ecology and the Environment, 14(7), 369–378. https://doi.org/10.1002/fee.1311

Kayanne, H., Harii, S., Ide, Y., & Akimoto, F. (2002). Recovery of coral populations after the 1998 bleaching on Shiraho Reef, in the southern Ryukyus, NW Pacific. Marine Ecology Progress Series, 239, 93–103. https://doi.org/10.3354/meps239093

Kayanne, H. (2017). Validation of degree heating weeks as a coral bleaching index in the northwestern Pacific. Coral Reefs, 36(1), 63–70. https://doi.org/10.1007/s00338-016-1524-y

Kelley, R. (2016). Indo Pacific Coral Finder, 3rd Edition. Townsville: BYOGUIDES.

Kushairi, M. R. M. (1998). The 1998 bleaching catastrophe of corals in the South China Sea. In Proc. JSPS Joint Seminar on Marine and Fisheries Sciences, Bali, Indonesia, 1998. http://ci.nii.ac.jp/naid/10018463620/en/

LaJeunesse, T. C., Pettay, D. T., Sampayo, E. M., Phongsuwan, N., Brown, B., Obura, D. O., et al. (2010). Long-standing environmental conditions, geographic isolation and host-symbiont specificity influence the relative ecological dominance and genetic diversification of coral endosymbionts in the genus Symbiodinium. Journal of Biogeography, 37(5), 785–800. https://doi.org/10.1111/j.1365-2699.2010.02273.x

Lenth, R. V. (2021). emmeans: Estimated Marginal Means, aka least-squares means. R package version 1.6.1.

Lesser, M. P. (1997). Oxidative stress causes coral bleaching during exposure to elevated temperatures. Coral reefs, 16(3), 187–192.

Liu, G., Heron, S. F., Mark Eakin, C., Muller-Karger, F. E., Vega-Rodriguez, M., Guild, L. S., et al. (2014). Reef-scale thermal stress monitoring of coral ecosystems: New 5-km global products from NOAA coral reef watch. Remote Sensing, 6(11), 11579–11606. https://doi.org/10.3390/rs61111579

Liu, G., Strong, A. E., & Skirving, W. (2003). Remote sensing of sea surface temperatures during 2002 Barrier Reef coral bleaching. Eos, Transactions American Geophysical Union, 84(15), 137–141. https://doi.org/10.1029/2003EO150001

Loya, Y., Sakai, K., Yamazato, K., Nakano, Y., Sambali, H., & Van Woesik, R. (2001). Coral bleaching: The winners and the losers. Ecology Letters, 4(2), 122–131. https://doi.org/10.1046/j.1461-0248.2001.00203.x

MacKellar, M. C., & McGowan, H. A. (2010). Air-sea energy exchanges measured by eddy covariance during a localised coral bleaching event, Heron Reef, Great Barrier Reef, Australia. Geophysical Research Letters, 37(24). https://doi.org/10.1029/2010GL045291

Marshall, P. A., & Baird, A. H. (2000). Bleaching of corals on the Great Barrier Reef: differential susceptibilities among taxa. Coral reefs, 19(2), 155–163.

Maynard, J. A., Turner, P. J., Anthony, K. R. N., Baird, A. H., Berkelmans, R., Eakin, C. M., et al. (2008). ReefTemp: An interactive monitoring system for coral bleaching using high-resolution SST and improved stress predictors. Geophysical Research Letters, 35(5), 1–5. https://doi.org/10.1029/2007GL032175

McClanahan, T. R., Ateweberhan, M., Graham, N. A. J., Wilson, S. K., Ruiz Sebastián, C., Guillaume, M. M. M., & Bruggemann, J. H. (2007a). Western Indian Ocean coral communities: Bleaching responses and susceptibility to extinction. Marine Ecology Progress Series, 337, 1–13. https://doi.org/10.3354/meps337001

McClanahan, T. R., Ateweberhan, M., Ruiz Sebastián, C., Graham, N. A. J., Wilson, S. K., Bruggemann, J. H., et al. (2007b). Predictability of coral bleaching from synoptic satellite and in situ temperature observations. Coral Reefs, 26(3), 695–701. https://doi.org/10.1007/s00338-006-0193-7

McClanahan, T. R., Baird, A. H., Marshall, P. A., & Toscano, M. A. (2004). Comparing bleaching and mortality responses of hard corals between southern Kenya and the Great Barrier Reef, Australia. Marine Pollution Bulletin, 48(3–4), 327–335. https://doi.org/10.1016/j.marpolbul.2003.08.024

McClanahan, T. R., & Maina, J. (2003). Response of coral assemblages to the interaction between natural temperature variation and rare warm-water events. Ecosystems, 6(6), 551– 56. https://doi.org/10.1007/s10021-002-0104-x

Morais, J., Morais, R. A., Tebbett, S. B., Pratchett, M. S., & Bellwood, D. R. (2021). Dangerous demographics in post-bleach corals reveal boom-bust versus protracted declines. Scientific Reports, 11(1), 1–7. https://doi.org/10.1038/s41598-021-98239-7

Muir, P. R., Marshall, P. A., Abdulla, A., & Aguirre, J. D. (2017). Species identity and depth predict bleaching severity in reef-building corals: Shall the deep inherit the reef? Proceedings of the Royal Society B: Biological Sciences, 284(1864). https://doi.org/10.1098/rspb.2017.1551

Muñiz-Castillo, A. I., & Arias-González, J. E. (2021). Drivers of coral bleaching in a Marine Protected Area of the Southern Gulf of Mexico during the 2015 event. Marine Pollution Bulletin, 166, 112256. https://doi.org/10.1016/j.marpolbul.2021.112256

Nakamura, T. V., & Van Woesik, R. (2001). Water-flow rates and passive diffusion partially explain differential survival of corals during the 1998 bleaching event. Marine Ecology Progress Series, 212, 301–304. doi: https://doi.org/10.3354/meps212301

Nelder, J. A., & Wedderburn, R. W. (1972). Generalized linear models. Journal of the Royal Statistical Society: Series A (General), 135(3), 370–384.

NOAA Coral Reef Watch Version 3.1 Daily Global 5-km Satellite Coral Bleaching Degree Heating Week Product. (2014). NOAA Coral Reef Watch Daily 5km Satellite CoralBleaching Heat Stress Monitoring Products (Version 3.1) [Accessed January 3, 2021]

Oliver, T. A., & Palumbi, S. R. (2011). Do fluctuating temperature environments elevate coral thermal tolerance? Coral Reefs, 30(2), 429–440. https://doi.org/10.1007/s00338-011-0721-y

Pachauri, R. K., Myles R. A., Vicente R. B., John B., Wolfgang C., Renate C., et al. (2014). Climate change 2014: synthesis report. Contribution of Working Groups I, II and III to the fifth assessment report of the Intergovernmental Panel on Climate Change (p. 151). Ipcc

Page, C. E., Leggat, W., Heron, S. F., Choukroun, S. M., Lloyd, J., & Ainsworth, T. D. (2019). Seeking resistance in coral reef ecosystems: the interplay of biophysical factors and bleaching resistance under a changing climate: the interplay of a reef’s biophysical factors can mitigate the coral bleaching response. BioEssays, 41(7), 1800226. https://doi.org/10.1002/bies.201800226

Palumbi, S. R., Barshis, D. J., Traylor-Knowles, N., & Bay, R. A. (2014). Mechanisms of reef coral resistance to future climate change. Science, 344(6186), 895–898. https://doi.org/10.1126/science.1251336

Pandolfi, J. M., Connolly, S. R., Marshall, D. J., & Cohen, A. L. (2011). Projecting coral reef futures under global warming and ocean acidification. Science, 333(6041), 418–422. http://paleodb.org/cgi-bin/bridge.pl

Paulay, G., & Benayahu, Y. (1999). Patterns and consequences of coral bleaching in Micronesia (Majuro and Guam) in 1992-1994. Micronesica-Agana, 32(1), 109–124. http://octocoralresearch.com/PDFFiles/Bleachinginoctocorals.pdf

Phongsuwan, N., & Chansang, H. (2012). Repeated coral bleaching in the Andaman Sea, Thailand, during the last two decades. Phuket Marine Biological Center Research Bulletin, 71, 19–41.

Pochon, X., Stat, M., Takabayashi, M., Chasqui, L., Chauka, L. J., Logan, D. D. K., & Gates, R. D. (2010). Comparison of endosymbiotic and free-living Symbiodinium (dinophyceae) diversity in a hawaiian reef environment. Journal of Phycology, 46(1), 53–65. https://doi.org/10.1111/j.1529-8817.2009.00797.x

Pratchett, M. S., McCowan, D., Maynard, J. A., & Heron, S. F. (2013). Changes in bleaching susceptibility among corals subject to ocean warming and recurrent bleaching in Moorea, French Polynesia. PLoS one, 8(7), e70443. https://doi.org/10.1371/journal.pone.0070443

Prosser, C. L. (Ed.). (1991). Comparative animal physiology, environmental and metabolic animal physiology. John Wiley & Sons.

Reef Check Malaysia. (2013). Status of Coral Reefs in Malaysia, 2013. http://www.reefcheck.org.my/media-information/annual-survey-reports

Reef Check Malaysia. (2019). Status of Coral Reefs in Malaysia, 2019. http://www.reefcheck.org.my/media-information/annual-survey-reports

Reef Check Malaysia. (2020). Status of Coral Reefs in Malaysia, 2020. http://www.reefcheck.org.my/media-information/annual-survey-reports

Richards, Z. T., Yasuda, N., Kikuchi, T., Foster, T., Mitsuyuki, C., Stat, M., et al. (2018). Integrated evidence reveals a new species in the ancient blue coral genus Heliopora (Octocorallia). Scientific Reports, 8(1), 1–14. https://doi.org/10.1038/s41598-018-32969-z

Roberts, T. E., Moloney, J. M., Sweatman, H. P. A., & Bridge, T. C. L. (2015). Benthic community composition on submerged reefs in the central Great Barrier Reef. Coral Reefs, 34(2), 569–580. https://doi.org/10.1007/s00338-015-1261-7

RStudio Team (2021). RStudio: Integrated Development Environment for R. RStudio, PBC, Boston, MA URL http://www.rstudio.com/. [Accessed April 1, 2021]

Safaie, A., Silbiger, N. J., McClanahan, T. R., Pawlak, G., Barshis, D. J., Hench, J. L., et al. (2018). High frequency temperature variability reduces the risk of coral bleaching. Nature Communications, 9(1), 1–12. https://doi.org/10.1038/s41467-018-04074-2

Schoepf, V., Grottoli, A. G., Levas, S. J., Aschaffenburg, M. D., Baumann, J. H., Matsui, Y., et al. (2015). Annual coral bleaching and the long-term recovery capacity of coral. Proceedings of the Royal Society B: Biological Sciences, 282 (1819). https://doi.org/10.1098/rspb.2015.1887

Schuhmacher, H., Loch, K., Loch, W., & See, W. R. (2005). The aftermath of coral bleaching on a Maldivian reef - A quantitative study. Facies, 51(1–4), 80–92. https://doi.org/10.1007/s10347-005-0020-6

Skirving, W. J., Heron, S. F., Marsh, B. L., Liu, G., De La Cour, J. L., Geiger, E. F., et al. (2019). The relentless march of mass coral bleaching: a global perspective of changing heat stress. Coral Reefs, 38(4), 547–557. https://doi.org/10.1007/s00338-019-01799-4

Stocker, T., Qin, D., Plattner, G., Alexander, L., Allen, S., Bindoff, N., et al (2014). Technical summary. In Climate change 2013: the physical science basis. Contribution of Working Group I to the Fifth Assessment Report of the Intergovernmental Panel on Climate Change (pp. 33–115). Cambridge University Press. https://doi.org/10.1017/cbo9781107415324.005

Strong, A. E., Liu, G., Kimura, T., Yamano, H., Tsuchiya, M., Kakuma, S. I., et al. (2002). Detecting and monitoring 2001 coral reef bleaching events in Ryukyu Islands, Japan using satellite bleaching HotSpot remote sensing technique. In IEEE International Geoscience and Remote Sensing Symposium (Vol. 1, pp. 237–239). IEEE. https://doi.org/10.1109/igarss.2002.1024998

Suggett, D. J., & Smith, D. J. (2020). Coral bleaching patterns are the outcome of complex biological and environmental networking. Global Change Biology, 26(1), 68–79. https://doi.org/10.1111/gcb.14871

Sully, S., Burkepile, D. E., Donovan, M. K., Hodgson, G., & van Woesik, R. (2019). A global analysis of coral bleaching over the past two decades. Nature Communications, 10(1), 1–5. https://doi.org/10.1038/s41467-019-09238-2

Tan, C. H., & Sf, H. (2011). First observed severe mass bleaching in Malaysia, Greater Coral Triangle. Galaxea, Journal of Coral Reef Studies, 13(1), 27–28.https://doi.org/10.3755/galaxea.13.27

Thomas, L., Rose, N. H., Bay, R. A., López, E. H., Morikawa, M. K., Ruiz-Jones, L., et al (2018). Mechanisms of thermal tolerance in reef-building corals across a fine-grained environmental mosaic: Lessons from Ofu, American Samoa. Frontiers in Marine Science, 4, 434. https://doi.org/10.3389/fmars.2017.00434

Thompson, D. M., & Van Woesik, R. (2009). Corals escape bleaching in regions that recently and historically experienced frequent thermal stress. Proceedings of the Royal Society B: Biological Sciences, 276(1669), 2893–2901. https://doi.org/10.1098/rspb.2009.0591

Toda, T., Okashita, T., Maekawa, T., Alfian, B. A. A. K., Rajuddin, M. K. M., Nakajima, R., Chen, W., Takahashi, K. T., Othman, B. H. R., & Terazaki, M. (2007). Community structures of coral reefs around peninsular Malaysia. Journal of Oceanography, 63(1), 113– 123. https://doi.org/10.1007/s10872-007-0009-6

Van Hooidonk, R., Maynard, J., Grimsditch, G., Williams, G., Tamelander, J., Gove, J., et al. (2020). Projections of future coral bleaching conditions using IPCC CMIP6 models: climate policy implications, management applications, and Regional Seas summaries. United Nation Environmental Programme, Nairobi, Kenya.

Van Hooidonk, R., Maynard, J., & Planes, S. (2013). Temporary refugia for coral reefs in a warming world. Nature Climate Change, 3(5), 508–511. https://doi.org/10.1038/nclimate1829

Van Hooidonk, R., Maynard, J., Tamelander, J., Gove, J., Ahmadia, G., Raymundo, L., et al. (2016). Local-scale projections of coral reef futures and implications of the Paris Agreement. Scientific Reports, 6(1), 1–8. https://doi.org/10.1038/srep39666

Van Hooidonk, R., Maynard, J., Tamelander, J., Gove, J., Ahmadia, G., Raymundo, L., et al. (2017). Coral Bleaching Futures: Downscaled Projections of Bleaching Conditions for the World’s Coral Reefs, Implications of Climate Policy and Management Responses. United Nation Environmental Programme, Nairobi, Kenya.

Van Woesik, R., Sakai, K., Ganase, A., & Loya, Y. (2011). Revisiting the winners and the losers a decade after coral bleaching. Marine Ecology Progress Series, 434, 67–76. https://doi.org/10.3354/meps09203

Van Woesik, R., Houk, P., Isechal, A. L., Idechong, J. W., Victor, S., & Golbuu, Y. (2012). Climate-change refugia in the sheltered bays of Palau: Analogs of future reefs. Ecology and Evolution, 2(10), 2474–2484. https://doi.org/10.1002/ece3.363

Voolstra, C. R., Buitrago-López, C., Perna, G., Cárdenas, A., Hume, B. C. C., Rädecker, N., et al. (2020). Standardized short-term acute heat stress assays resolve historical differences in coral thermotolerance across microhabitat reef sites. Global Change Biology, 26 (8), 4328– 4343. https://doi.org/10.1111/gcb.15148

Voolstra, C. R., Valenzuela, J. J., Turkarslan, S., Cárdenas, A., Hume, B. C. C., Perna, G., et al. (2021). Contrasting heat stress response patterns of coral holobionts across the Red Sea suggest distinct mechanisms of thermal tolerance. Molecular Ecology, 30(18), 4466–4480. https://doi.org/10.1111/mec.16064

Wellington, G. M., Glynn, P. W., Strong, A. E., Navarrete, S. A., Wieters, E., & Hubbard, D. (2001). Crisis on coral reefs linked to climate change. Eos, Transactions American Geophysical Union, 82(1), 1–5.

Wong, K. H., Goodbody-Gringley, G., de Putron, S. J., Becker, D. M., Chequer, A., & Putnam, H. M. (2021). Brooded coral offspring physiology depends on the combined effects of parental press and pulse thermal history. Global Change Biology, 27(13), 3179–3195. https://doi.org/10.1111/gcb.15629

Wooldridge, S. A., Done, T. J., Thomas, C. R., Gordon, I. I., Marshall, P. A., & Jones, R. N. (2012). Safeguarding coastal coral communities on the central Great Barrier Reef (Australia) against climate change: Realizable local and global actions. Climatic Change, 112(3–4), 945–961. https://doi.org/10.1007/s10584-011-0229-z

Zann, L. P., & Bolton, L. (1985). The distribution, abundance and ecology of the blue coral Heliopora coerulea (Pallas) in the Pacific. Coral Reefs, 4(2), 125–134. https://doi.org/10.1007/BF0030087

Zvuloni, A., Artzy-Randrup, Y., Stone, L., van Woesik, R., & Loya, Y. (2008). Ecological size-frequency distributions: how to prevent and correct biases in spatial sampling. Limnology and Oceanography: Methods, 6(3), 144–153.

