## Supplementary materials for "Highly variable response of hard coral taxa to successive coral bleaching events (2019-2020) and rising ocean temperatures in Northeast Peninsular Malaysia"

**Supplementary S1|** List of anthozoan taxa observed bleached off-transect during surveys in June 2019 in Pulau Lang Tengah in Northeast Peninsular Malaysia.

| **Non-scleractinian** | **Common name** | **Species name** |
|---|---|---|
| Anemone | Magnificent Anemone | *Heteractis magnifica; Heteractis crispa* |
| Anemone | Bubble Tip Anemone | *Entacmaea quadricolor* |
| Anemone | Carpet Anemone | *Stichodactyla gigantea* |
| Fire Coral | - | *Millepora cataphyllia* |
| Soft Coral | - | *Sarcophyton spp.* |
| Soft Coral | - | *Sinularia spp.* |
| Zoanthid | - | *Palythoa tuberculosa* |
| **Scleractinian** | **Colony growth form** | **Species name** |
| *Alveopora* | Submassive | *A. allingi* |
| *Alveopora* | Submassive | *A. catalai* |
| *Alveopora* | Submassive | *A. gigas* |
| *Astrea* | Submassive | *A. curta* |
| *Astrea* | Submassive | *A. annuligrea* |
| *Cycloseris* | Solitary | *Cycloseris spp.* |
| *Ctenactis* | Solitary | *C. albitentaculata* |
| *Ctenactis* | Solitary | *C. crassa* |
| *Ctenactis* | Solitary | *C. echinata* |
| *Gardineroseris* | Submassive | *G. Planulata* |
| *Leptoseris* | Encrusting (plate) | *L. scabra* |
| *Lithophyllon* | Encrusting (plate) | *L. undulatum* |
| *Montipora* | Encrusing with upgrowth | *M. hispida* |
| *Phymastrea* | Submassive | *P. valenciennesi* |
| *Porites* | Branching | *P. cylindrica* |
| *Psammocora* | Columnar | *P. digitata* |

**Supplementary S2|** Mean annual sea surface temperature (SST°C) and mean bleaching season (April-September) sea surface temperatures of the 10 warmest years since 1985 are shown for Pulau Lang Tengah, Northeast Peninsular Malaysia. SST was recorded by the National Oceanic and Atmospheric Association (NOAA), Coral Reef Watch (CRW) product, version 3.1, 2014.

| Year | Annual mean SST (°C) | Year | Bleaching season mean SST (°C) |
|---|---|---|---|
| 2010 | 29.65 | 2010 | 30.38 |
| 2016 | 29.51 | 2020 | 30.24 |
| 1998 | 29.48 | 1998 | 30.13 |
| 2020 | 29.47 | 2019 | 30.12 |
| 2019 | 29.45 | 2014 | 30.1 |
| 2017 | 29.39 | 2016 | 30.09 |
| 2018 | 29.25 | 2017 | 30.02 |
| 2013 | 29.19 | 2013 | 29.97 |
| 2014 | 29.19 | 2006 | 29.93 |
| 2012 | 29.18 | 2009 | 29.89 |

**Supplementary S3|** Summary of recorded thermal stress (DHW °C –weeks) in 2019 and 2020 at numerous locations across the eastern coast of Peninsular Malaysia. Heat stress was recorded by the National Oceanic and Atmospheric Association (NOAA), Coral Reef Watch (CRW) product, version 3.1, 2014.

| Location | Coordinates | DHW 2019 | DHW 2020 | Peak DHW 2019 | Peak DHW 2020 |
|---|---|---|---|---|---|
| Pulau Perhentian | 5°56'02.5"N,102°43'51.5"E | 2.00 | 0.15 | 01.06.2019 | 31.05.2020 |
| Pulau Lang Tengah | 5°46'30.0"N, 102°52'30.0"E | 1.05 | 0.00 | 01.06.2019 | - |
| Pulau Yu Kecil | 5°37'28.3"N, 103°09'38.4"E | 2.53 | 0.00 | 01.06.2019 | - |
| Pulau Tenggol | 4°48'14.1"N, 103°40'46.5"E | 2.65 | 0.47 | 12.05.2019 | 22.05.2020 |
| Pulau Tioman | 2°47'58.6"N, 104°10'09.5"E | 1.74 | 0.49 | 09.05.2019 | 22.05.2020 |
| Pulau Sibu | 2°13'14.7"N, 104°03'16.7"E | 4.61 | 1.90 | 15.05.2019 | 21.05.2020 |

**Supplementary S4|** High-frequency diel temperature variations are presented as average daily (24h) temperature ranges (DTR) recorded at Pulau Lang Tengah at three logging stations at 8 meters of water depth. Here, House Reef (HR) is leeward, Batu Kucing 1 (BK1) and Tanjung Telunjuk 1 (TT1) are windward. All loggers recorded at an interval of 90 minutes. DTR time series include: ($DTR_{Total}$: September 2019 to September 2020), bleaching season ($DTR_{BS}$: April to September 2020), peak summer season ($DTR_{SS}$: May-July 2020), monsoon season in boreal fall and winter ($DTR_{FW}$: October-March), and for the 60 and 90 day period preceding the first bleaching observation in 2020 (June 20th, 2020), respectively ($DTR_{60}$ and $DTR_{90}$). A non-parametric Kruskal-Wallis test was done to investigate seasonal differences in DTR distribution across sites at significance levels of *** p<0.001, **p< 0.01, *p< 0.05, ns – not significant.

| | Site | | | Kruskal-Wallis | |
|---|---|---|---|---|---|
| | HR | BK 1 | TT 1 | H | p |
| $DTR_{Total}$ | 0.55 | 0.44 | 0.48 | 81.32 | *** |
| $DTR_{BS}$ | 0.65 | 0.45 | 0.53 | 43.10 | *** |
| $DTR_{SS}$ | 0.61 | 0.39 | 0.49 | 57.62 | *** |
| $DTR_{FW}$ | 0.43 | 0.39 | 0.41 | 1.18 | ns |
| $DTR_{60}$ | 0.59 | 0.41 | 0.50 | 19.93 | *** |
| $DTR_{90}$ | 0.58 | 0.41 | 0.50 | 7.91 | * |

**Supplementary S5|** General table showing the taxon-specific bleaching resistance (BRI) and bleaching induced mortality index (BIMI) during successive marine heatwaves in June 2019 and June 2020 in Pulau Lang Tengah, Northeast Peninsular Malaysia. The percentage of partial and whole-colony mortality of scleractinian taxa is shown for October 2019 to highlight post-bleaching mortality after heat disturbance in June 2019.

| | | June 2019 | | October 2019 | | | June 2020 | |
|---|---|---|---|---|---|---|---|---|
| ***Genus*** | **Morphology** | **n (colonies)** | **BRI** | **n (colonies)** | **Dead (%)** | **BIMI** | **n (colonies)** | **BRI** |
| *Acropora* | Arborescent | 26 | 88 | 17 | 5.9 | 1.47 | 15 | 100 |
| | Corymbose | 32 | 95 | 25 | 0.0 | 0.00 | 33 | 92 |
| | Digitate | 9 | 91 | 12 | 0.0 | 0.00 | 11 | 100 |
| | Hispidose | 143 | 74 | 100 | 0.0 | 0.00 | 102 | 100 |
| | Tabular | 10 | 86 | 4 | 25.0 | 12.50 | 11 | 100 |
| *Astreopora* | Massive | 1 | 60 | 1 | 0.0 | 0.00 | 1 | 60 |
| | Encrusting | 1 | 0 | 1 | 0.0 | 0.00 | 0 | 0 |
| *Blasstomussa* | Encrusting | 0 | 0 | 1 | 0.0 | 0.00 | 3 | 100 |
| *Cyphastrea* | Encrusting | 19 | 40 | 25 | 0.0 | 0.00 | 22 | 92 |
| *Diploastrea* | Encrusting | 2 | 90 | 2 | 0.0 | 0.00 | 2 | 90 |
| *Echinopora* | Branching | 19 | 14 | 11 | 0.0 | 0.00 | 7 | 86 |
| | Encrusting | 15 | 83 | 14 | 7.1 | 3.57 | 17 | 100 |
| *Favia* | Encrusting | 11 | 100 | 17 | 0.0 | 0.00 | 31 | 97 |
| | Submassive | 47 | 76 | 74 | 2.7 | 0.68 | 76 | 96 |

| | | | | | | | | |
|---|---|---|---|---|---|---|---|---|
| | Massive | 3 | 47 | 5 | 40.0 | 15.00 | 7 | 83 |
| *Favites* | Encrusting | 48 | 57 | 35 | 2.9 | 0.71 | 35 | 93 |
| *Fungia* | Solitary | 115 | 61 | 120 | 0.8 | 0.42 | 137 | 91 |
| *Galaxea* | Encrusting | 96 | 98 | 110 | 0.0 | 0.00 | 101 | 95 |
| *Goniastrea* | Submassive | 52 | 34 | 61 | 0.0 | 0.00 | 44 | 85 |
| *Goniopora* | Submassive | 0 | 0 | 4 | 0.0 | 0.00 | 3 | 100 |
| *Herpolitha* | Solitary | 2 | 40 | 4 | 0.0 | 0.00 | 1 | 100 |
| *Heliopora* | Columnar | 122 | 18 | 99 | 14.1 | 11.36 | 92 | 59 |
| *Hydnophora* | Branching | 1 | 60 | 1 | 0.0 | 0.00 | 1 | 100 |
| | Encrusting | 6 | 60 | 7 | 0.0 | 0.00 | 6 | 90 |
| *Leptastrea* | Encrusting | 69 | 93 | 72 | 0.0 | 0.00 | 80 | 92 |
| *Leptoria* | Submassive | 2 | 100 | 1 | 0.0 | 0.00 | 1 | 100 |
| *Merulina* | Encrusting | 7 | 69 | 6 | 0.0 | 0.00 | 5 | 100 |
| *Montipora* | Encrusting | 15 | 71 | 8 | 0.0 | 0.00 | 7 | 91 |
| | Vase | 36 | 86 | 32 | 6.3 | 3.91 | 37 | 97 |
| *Oulophyllia* | Massive | 5 | 88 | 2 | 0.0 | 0.00 | 3 | 100 |
| *Pachyseris* | Encrusting | 3 | 33 | 2 | 0.0 | 0.00 | 1 | 100 |
| *Pavona* | Encrusting | 9 | 100 | 11 | 0.0 | 0.00 | 7 | 100 |
| | Foliose | 135 | 37 | 114 | 4.4 | 1.97 | 123 | 73 |
| *Physogyra* | Submassive | 1 | 100 | 7 | 0.0 | 0.00 | 5 | 100 |
| *Platygyra* | Encrusting | 12 | 90 | 17 | 5.9 | 1.47 | 23 | 84 |
| | Submassive | 56 | 69 | 78 | 2.6 | 1.60 | 80 | 81 |
| *Pocillopora* | Corymbose | 29 | 60 | 31 | 6.5 | 4.84 | 38 | 85 |
| *Porites* | Encrusting | 47 | 48 | 68 | 1.5 | 1.47 | 76 | 90 |
| *Porites sp. 2* | Encrusting | 447 | 66 | 384 | 1.3 | 3.20 | 354 | 88 |
| *Porites* | Massive | 155 | 47 | 164 | 6.7 | 0.78 | 189 | 79 |
| *Psammocora* | Encrusting | 18 | 97 | 16 | 0.0 | 0.00 | 22 | 100 |
| | Foliose | 24 | 99 | 29 | 0.0 | 0.00 | 26 | 100 |
| *Sandolitha* | Solitary | 1 | 80 | 3 | 0.0 | 0.00 | 3 | 100 |
| *Stylocoenellia* | Cryptic | 5 | 100 | 0 | 0.0 | 0.00 | 6 | 100 |
| *Symphyllia* | Submassive | 26 | 94 | 31 | 3.2 | 3.23 | 34 | 96 |
| **Total** | | **1882** | **64** | **1826** | **2.9** | **1.71** | **1884** | **88** |

**Supplementary S6|** Comparison of bleaching incidence and mortality of hard corals in Pulau Tioman in 2010 [as recorded by Guest et al. (2012)] and Pulau Lang Tenagh in 2019. Mortality incidence for Lang Tengah Island was recorded in October 2019. For surveys in Pulau Lang Tengah, moderate bleaching includes categories pale live and fluorescent (B2), and colonies with ≤66% of surface area bleached (B3 and B4). Severe bleaching includes all colonies with >66% of the colony surface area bleached (categories B5-B6).

| | Pulau Tioman 2010 (Guest et al. 2012) | | | | | Pulau Lang Tengah 2019 | | | | |
|---|---|---|---|---|---|---|---|---|---|---|
| **Genus** | **n** | **Normal (%)** | **Moderate (%)** | **Severe (%)** | **Dead (%)** | **n** | **Normal (%)** | **Moderate (%)** | **Severe (%)** | **Dead (%)** |
| *Acropora* | 632 | 70 | 3 | 1 | 26 | 220 | 30 | 67 | 3 | 1 |
| *Pocillopora* | 87 | 39 | 21 | 2 | 38 | 29 | 38 | 28 | 34 | 7 |
| *Hydnophora* | 32 | 3 | 3 | 41 | 53 | 7 | 29 | 43 | 29 | 0 |
| *Echinopora* | 8 | 100 | 0 | 0 | 0 | 34 | 35 | 12 | 53 | 4 |
| *Favites* | 53 | 91 | 9 | 0 | 0 | 48 | 37 | 29 | 34 | 3 |
| *Symphyllia* | 7 | 86 | 14 | 0 | 0 | 26 | 69 | 31 | 0 | 0 |
| *Platygyra* | 43 | 86 | 9 | 2 | 2 | 68 | 25 | 62 | 13 | 0 |
| *Galaxea* | 56 | 91 | 7 | 2 | 0 | 96 | 92 | 8 | 0 | 0 |
| *Favia* | 19 | 100 | 0 | 0 | 0 | 61 | 66 | 18 | 16 | 0 |
| *Fungia* | 128 | 54 | 38 | 5 | 4 | 115 | 17 | 58 | 25 | 1 |
| *Pavona* | 6 | 17 | 0 | 67 | 17 | 144 | 38 | 5 | 57 | 4 |
| *Goniastrea* | 71 | 90 | 6 | 0 | 4 | 52 | 29 | 6 | 65 | 0 |
| *Cyphastrea* | 15 | 87 | 13 | 0 | 0 | 19 | 32 | 16 | 53 | 0 |
| *Montipora* | 418 | 78 | 4 | 0 | 17 | 51 | 65 | 24 | 12 | 5 |
| *Porites 1 +2* | 86 | 74 | 23 | 2 | 0 | 649 | 46 | 18 | 36 | 3 |

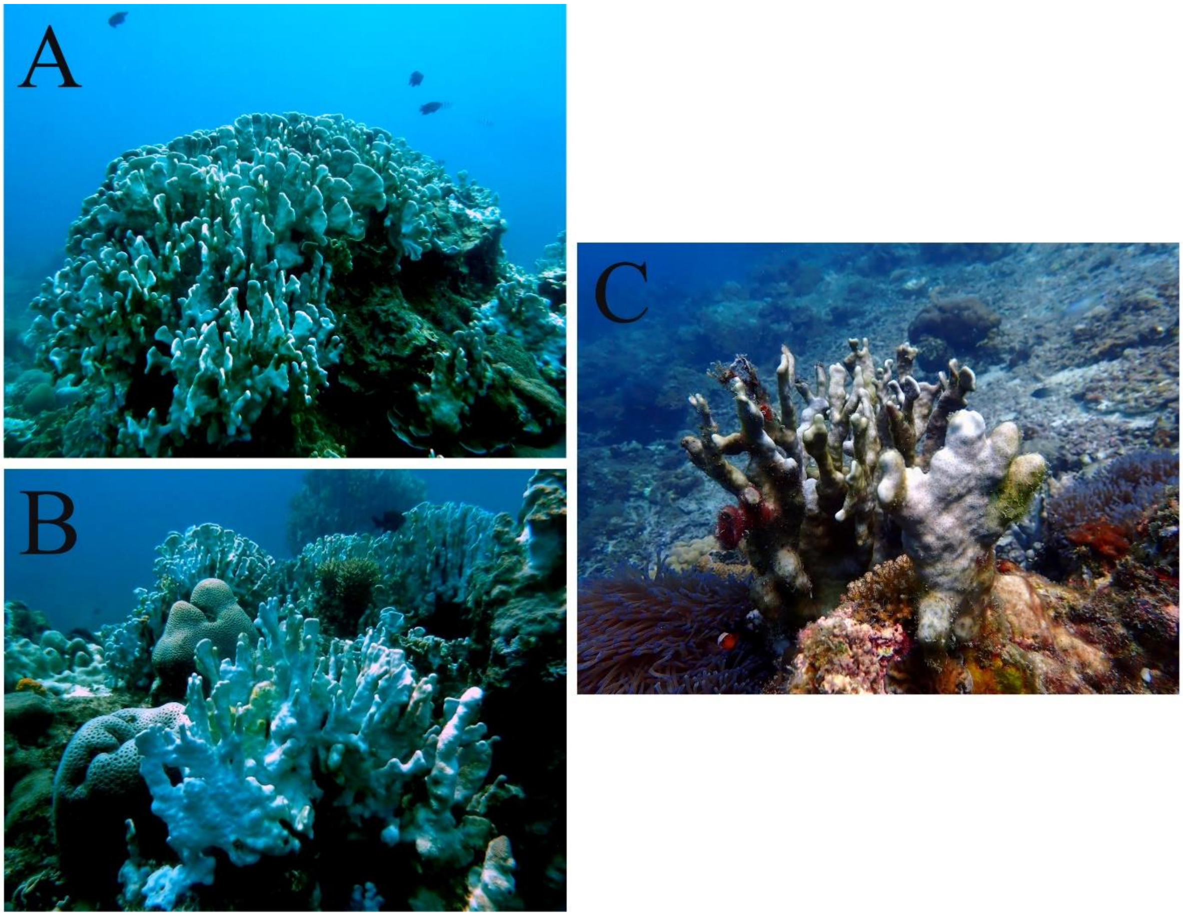


**Supplementary Figure S7|** High bleaching incidence and severity of *Heliopora sp*. colonies was recorded in June 2019 and June 2020 around Pulau Lang Tengah in Northeast Peninsular Malaysia: (A) shows a large *Heliopora sp*. colony on a shallow (5-6 m) fringing reef; (B) highlights high bleaching incidence of *Heliopora sp*. at windward sites and subsequent partial mortality of individual colonies (C).
